# Domain-Specific Agonist Binding Affinities Explain Structural and Functional Regulation of TRPM2

**DOI:** 10.64898/2026.03.30.715250

**Authors:** Tatiana Kupriianova, Tessa Schwarzer, Thorben Thalacker, Lucas Defelipe, Stefanie Etzold, Frederike Kulow, Vanessa Pahl, Shivansh Goyal, Vanna Nguyen, Michael Zimmermann, Andreas H. Guse, Xiaolu A. Cambronne, Henning Tidow, Ralf Fliegert, Maria Garcia Alai

**Affiliations:** European Molecular Biology Laboratory, Hamburg, Germany; Centre for Structural Systems Biology, Hamburg, Germany; The Calcium Signalling Group, Department of Biochemistry and Molecular Cell Biology, University Medical Center Hamburg-Eppendorf, Martinistrasse 52, 20246 Hamburg, Germany; Hamburg Advanced Research Centre for Bioorganic Chemistry (HARBOR) & Department of Chemistry, Institute for Biochemistry and Molecular Biology, University of Hamburg, Luruper Chaussee 149, 22761 Hamburg, Germany; European Molecular Biology Laboratory, Heidelberg, Germany; Department of Molecular Biosciences, University of Texas at Austin, Austin, TX, USA

## Abstract

TRPM2 is a Ca²⁺-permeable cation channel activated by ADP-ribose (ADPR) and oxidative stress, yet the relative contributions of its two nucleotide-binding domains, MHR1/2 and NUDT9H, remain incompletely understood. Here, we quantitatively determine the affinities of the isolated human TRPM2 MHR1/2 and NUDT9H domains for ADPR, 2’-deoxy-dADPR (dADPR), and 8-Br-cADPR using biophysical approaches. The MHR1/2 domain binds ADPR with high affinity (K_d_ ≈ 0.5 µM), whereas the NUDT9H domain displays substantially lower affinity (K_d_ ≈ 192 µM), revealing a difference of nearly three orders of magnitude. Mutational analysis demonstrates that alterations in MHR1/2 strongly affect ligand binding and channel activation, while mutations within NUDT9H that markedly reduce ligand affinity exert only modest effects on gating. In parallel, we quantify intracellular ADPR concentrations in resting and hydrogen peroxide–stimulated cells and find that they remain well below the affinity required for substantial NUDT9H occupancy. Together, our findings indicate that high-affinity binding to the MHR1/2 domain is sufficient to drive TRPM2 activation under physiological conditions, whereas the NUDT9H domain likely contributes to maintaining the structural integrity of the channel rather than directly mediating ligand-dependent activation. These results provide a quantitative framework for understanding ligand-dependent TRPM2 regulation in cells.

## Introduction

Oxygen is often described as a *Janus gas*: it is essential for life, yet it can also cause harm. While oxygen fuels ATP production in cells, it also generates reactive oxygen species (ROS)^1^. At controlled levels, ROS participate in normal cell signaling^2^. However, when ROS accumulate, they damage proteins, lipids, and DNA, and drive processes such as inflammation, cell death, and neurodegeneration^3,4^. Cells have therefore evolved sensitive molecular sensors to detect redox imbalance and translate oxidative stress into adaptive responses. Among these sensors, ion channels occupy a pivotal position, coupling oxygen-induced stress triggers to the universal language of Ca²⁺ signaling.

Transient receptor potential melastatin 2 (TRPM2) is one of these channels (Figure 1A). As a nonselective cation channel permeable to Ca²⁺, TRPM2 transforms cues of oxidative stress into Ca²⁺ signals that shape immune activation, inflammation, and cell death^5,6^. As oxidative stress and inflammation play a key role in the development of a wide variety of diseases, TRPM2 serves as a central hub in conditions ranging from ischemia–reperfusion injury and stroke to diabetes and neurodegeneration^7^. Therefore, TRPM2 is an attractive pharmacological target and investigating this protein could reveal novel therapeutic strategies for treating these disorders.

**Figure 1.**
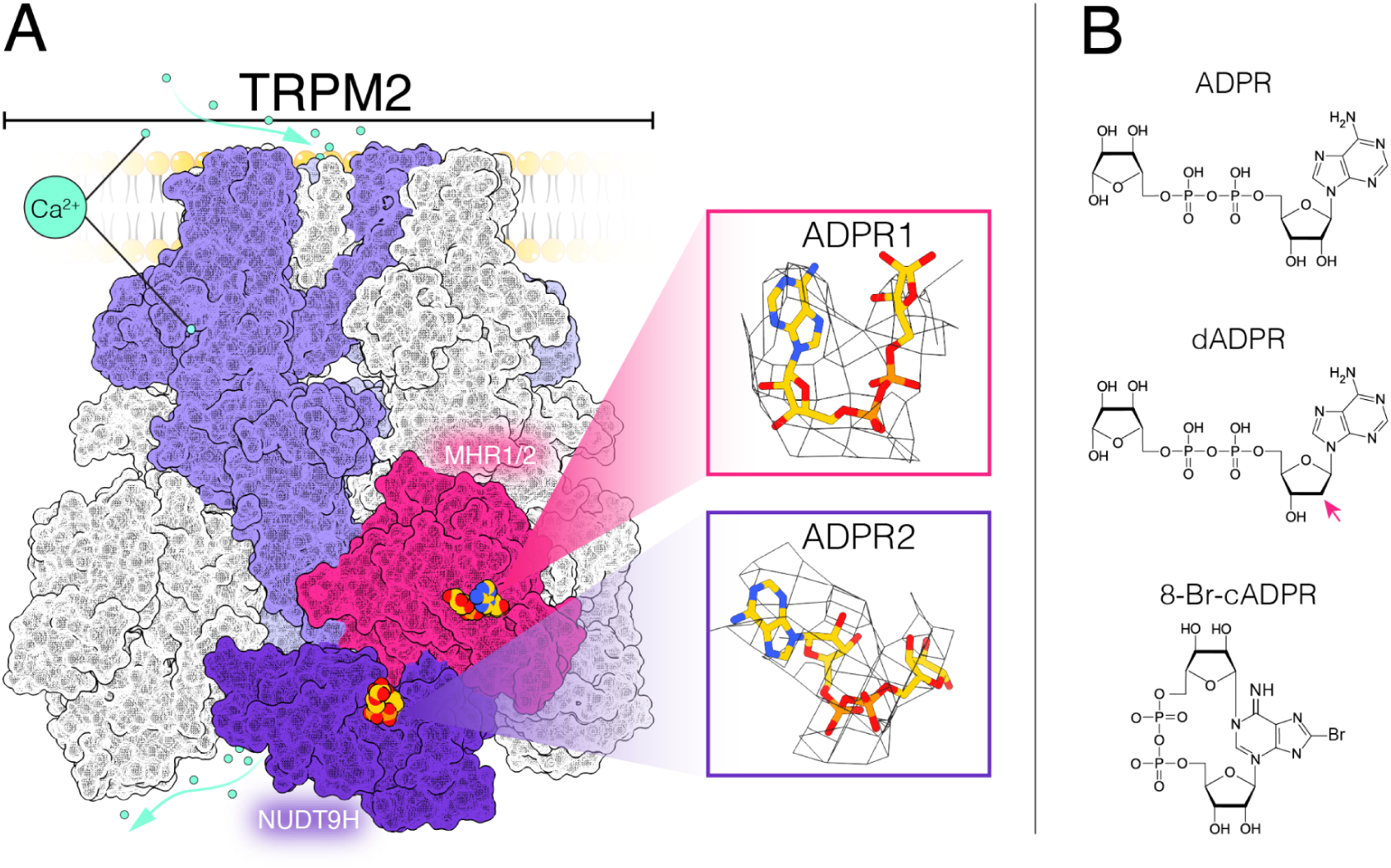
Overview of the calcium channel TRPM2 and its ligands. (A) Schematic surface representation of the human TRPM2 Cryo-EM structure (PDB IDs: 6PUS)^16^. Different subunits are shown in white and lilac. On a single subunit, the MHR1/2 domain is highlighted in magenta and the NUDT9H domain in violet. TRPM2 is bound to ligands: the agonist Ca²⁺ (green-cyan) in the transmembrane domain and ADPR (yellow) in the MHR1/2 and NUDT9H binding pockets. Ligands are shown on top of the surface representation to indicate their approximate binding sites. (B) Chemical structures of TRPM2 ligands ADPR, dADPR, and 8-Br-cADPR. The arrow highlights the difference between ADPR and dADPR - absence of oxygen at the 2′ position.

At the heart of TRPM2’s response to oxidative stress is its sensitivity to ADP-ribose (ADPR) (Figure 1B). ADPR is produced during oxidative stress, DNA repair, and NAD⁺ metabolism. A major producer of ADPR is the poly-(ADP-ribose) polymerase (PARP)/ Poly(ADP-ribose) glycohydrolase (PARG) system, which becomes highly active under oxidative conditions^8^. Hydrogen peroxide (H₂O₂) causes DNA strand breaks that activates PARP-1/2. These enzymes consume NAD⁺ to generate long chains of poly(ADP-ribose) (PAR) on nuclear proteins. PARG then degrades these PAR chains, releasing free monomeric ADPR which can leave the nucleus via nuclear pores. In parallel, CD38 - a NAD⁺-consuming type II ectoenzyme, a small fraction of which is also present as a type III transmembrane protein with its active site facing the cytoplasm - also generates ADPR during immune activation and inflammation^9^. Together, these pathways ensure that ROS and inflammatory signaling converge on the production of ADPR, which directly activates TRPM2.

Besides ADPR, 2′-deoxy-ADPR (dADPR) has been identified as a more efficacious endogenous “superagonist” (Figure 1B)^10^. dADPR differs from ADPR by missing just one oxygen atom, but the effect is dramatic: dADPR is about ten times more efficacious, activates the channel much faster, and works even at very low Ca²⁺ levels, where ADPR shows only minimal activity^10,11^. dADPR can be produced in cells through the pathway: NMN + dATP → dNAD → (via CD38) → dADPR. Its levels can be influenced by oxidative stress. Oxidative stress halts DNA replication and causes dATP to accumulate. The elevated dATP then favors its conversion into dADPR, resulting in activation of TRPM2 channels. This mechanism directly links cellular stress to TRPM2 signaling. Unlike ADPR, whose production consumes NAD, dADPR can activate TRPM2 without depleting this essential metabolite, and that is why it may serve as a signal for physiological processes rather than for cell death^10^.

The search for nucleotide binding sites in TRPM2 has uncovered a fascinating evolutionary story. At its C-terminus, TRPM2 carries the NUDT9 homology (NUDT9H) domain, which is homologous to the mitochondrial ADPR-pyrophosphatase NUDT9^12^. This homology initially led Perraud and coworkers to propose that the NUDT9H domain functions both as an ADPR binding site and an ADPR-hydrolyzing enzyme^12^. However, a look at the evolution of the channel reveals a more complex picture: sometime between early chordates and vertebrates, the NUDT9H domain gradually lost its enzymatic activity^13^. In invertebrates, such as *Nematostella vectensis*, the NUDT9H domain retains its ability to hydrolyze ADPR. Surprisingly, mutations that abolish the enzymatic activity of the NUDT9H domain make *nv*TRPM2 sensitive to oxidative stress^14^. Moreover, complete removal of the NUDT9H domain in *nv*TRPM2 does not interfere with activation of the channel by ADPR, suggesting that additional nucleotide binding sites contribute to channel regulation^14^.

Subsequent cryo-EM studies shed light on the mechanism of TRPM2 activation by identifying a second ADPR binding site in the N-terminal TRPM homology region (MHR1/2)^15–17^. Despite the growing structural insight, the functional interplay between the MHR1/2 domain and the NUDT9H domain remains elusive, particularly in vertebrate TRPM2 orthologues. While in the earlier TRPM2 orthologues the catalytic NUDT9H domain is weakly attached to the residual channel, in vertebrate TRPM2, the NUDT9H appears to be more tightly integrated with the remaining part of the channel. Removal of the NUDT9H domain in *hs*TRPM2 and *dr*TRPM2 completely abrogates activation by TRPM2, and individual point mutations in the NUDT9H domain thought to interfere with ADPR binding, did either abrogate or reduce the current induced by the agonist. Interestingly, also cryoEM indicates different domain interactions for *hs*TRPM2 and *dr*TRPM2, the NUDT9H domains can be exchanged between the orthologues without losing activation by ADPR^18^. Furthermore, a recent preprint indicates that an isolated NUDT9H domain of *hs*TRPM2, when expressed as an independent protein, can integrate with *hs*TRPM2 ΔNUDT9H thereby restoring gating by ADPR^19^. For the pre-vertebrate *sr*TRPM2 it was shown that tetramerization of the semi-independent NUDT9H domain contributes to the rotation of the MHR3/4 subdomain, which opens the ion conduction pathway thereby modulating gating kinetics^20^. Huang et al. propose that, in vertebrate channels the NUDT9H domain has become more tightly integrated with the rest of the channel. It forms a gating ring that couples ADPR binding to conformational changes that propagate to the pore, making it part of the activation machinery.

In addition, the role of the separate binding site has been explored using site specific modulators. For *hs*TRPM2 the cADPR antagonist 8-Br-cADPR^21^ has been shown to bind exclusively to the MHR1/2 nucleotide-binding pocket (Figure 1A)^16^, the selectivity being considered a consequence of the different shape of the binding pockets: ADPR binds to the NUDT9H domain in an elongated conformation whereas it adopts a U-shaped conformation in the MHR1/2 domain. The cyclic form of 8-Br-cADPR mimics this U-shape, which might allow it to occupy the MHR1/2 binding site but not the NUDT9H binding site. The bromide group would clash with Y295, preventing full insertion and stabilising TRPM2 in an inactive, apo-like conformation, effectively blocking the channel^16^. Of note, we found recently that the MHR1/2 domain of *dr*TRPM2 does not interact with the cyclic nucleotide while binding linear 8-Br-ADPR with high affinity^22^.

Inosine 5′-diphosporibose (IDPR), a substrate of the enzyme NUDT9, has been shown to activate both *nv*TRPM2 and *hs*TRPM2 but was in contrast to ADPR not capable to activate nvTRPM2 ΔNUDT9H, indicating that IDPR acts selectively on the nucleotide binding site in the NUDT9H domain. Importantly there is currently little biophysical data regarding the affinity of the two ADPR binding domains for the modulating nucleotides: Two earlier studies provided largely divergent K_d_ values for binding of ADPR to the NUDT9H domain of *hs*TRPM2 (K_d_ = 130 µM determined by ITC^23^, K_d_ ∼15 µM determined by SPR^15^), but no information is available for the K_d_ of the MHR1/2 domain of *hs*TRPM2 and for the nucleotides dADPR and 8-Br-cADPR.

In the present study, we address this gap by quantitatively determining the affinities of the isolated MHR1/2 and NUDT9H domains for ADPR, dADPR, and 8-Br-cADPR using complementary biophysical approaches. We further assess the impact of selected point mutations on ligand binding to these domains and evaluate their functional relevance. In parallel, we determine intracellular ADPR concentrations in unstimulated cells and following oxidative stress induced by H_2_O_2_. By integrating biochemical, structural, and cellular measurements, our work provides a quantitative framework for understanding how ligand binding and domain-specific interactions contribute to TRPM2 activation under physiological conditions.

## Results

### ADPR-binding sites in human TRPM2 bind their ligands with different affinities

Using nano-differential scanning fluorimetry (nDSF)^24^ we measured the binding affinities (K_d_) for ADPR and dADPR for each domain using separately expressed human NUDT9H and MHR1/2 constructs (Supl. Figure 1). The MHR1/2 domain exhibited much higher binding affinity for both ADPR and dADPR, with K_d_ values with asymmetric 95% confidence intervals (CI95) 0.51 µM [0.44 µM; 0.66 µM] and 0.21 µM, [0.17 µM; 0.32 µM] respectively (Supl. Figure 1). In contrast, the NUDT9H domain bound these ligands with three orders of magnitude lower affinity (K_d_ values with CI95 are 192 µM [160 µM; 220 µM] and 124 µM [110 µM; 140 µM], respectively). The results for the NUDT9H domain were further confirmed by isothermal titration calorimetry (ITC), which yielded similar low affinities (K_d_ = 167 µM for ADPR and 165 µM for dADPR). These K_d_ values for the NUDT9H domain are similar to the K_d_ = 130 µM previously determined by Grubisha and coworkers using ITC, but higher than the K_d_ of 15 µM for the isolated NUDT9H domain^25^ and 41 µM for the full *hs*TRPM2 reported by Wang and coworkers using surface plasmon resonance (SPR)^15^.

For both domains there was no significant difference between ADPR and dADPR in terms of binding affinity (Supl. Figure 1). This is in line with our previous finding that ADPR and dADPR do not differ in potency^10^, but that dADPR opens TRPM2 faster, indicating its enhanced effect is due to gating or conformational changes rather than higher affinity^11^.

### 8-Br-cADPR rapidly degrades under experimental conditions

In addition to ADPR and dADPR, we also investigated the interaction of the MHR1/2 and NUDT9H domain with 8-Br-cADPR. Interestingly, we observed binding of 8-Br-cADPR to the NUDT9H domain, despite previous structural data indicating that 8-Br-cADPR interacts selectively with the MHR1/2 domain (Supl. Figure 2)^14^. We hypothesized that this apparent discrepancy might result from degradation of 8-Br-cADPR into linear 8-Br-ADPR, which could in turn interact with the NUDT9H domain. To test this, we analyzed the stability of 8-Br-cADPR by HPLC (Supl. Figure 3), which revealed degradation of 8-Br-cADPR, not only into 8-Br-ADPR but also into cADPR and ADPR. Control samples, even without heating, showed some spontaneous breakdown: 8-Br-ADPR converted to ADPR, while 8-Br-cADPR degraded to cADPR. Under nDSF conditions (heating up to 70 °C in the presence of TCEP), 8-Br-ADPR further degraded into ADPR, whereas 8-Br-cADPR predominantly yielded cADPR, along with detectable amounts of 8-Br-ADPR and ADPR. Importantly, these degradation patterns were independent of the protein present during incubation (Supl. Figure 3).

Given that the assay buffer contained only 50 mM TrisHCl and 0.5 mM TCEP, we considered either the reducing agent or elevated temperature as potential drivers of degradation. We therefore compared the stability of 8-Br-cADPR and 8-Br-ADPR at room temperature and 50 °C, with or without TCEP (Supl. Figure 4). These experiments demonstrated that TCEP or heat alone had only minor effects, whereas their combination strongly promoted breakdown of the nucleotide. Previous studies have shown that TCEP is capable of removing bromine from organic compounds, e.g. it can reductively dehalogenate monobromobimane (MBB)^26^. In this reaction, the trivalent phosphine attacks the brominated carbon, releasing bromide and forming a reduced product. This supports our observation that the 8-brominated nucleotides become dehalogenated in our experiments.

Together, these results suggest that 8-brominated nucleotides are unstable under nDSF conditions because TCEP and elevated temperature synergistically accelerate their degradation. Consequently, nDSF is not suitable for reliably analysing the binding of 8-brominated nucleotides.

### Key residues involved in binding of ADPR and dADPR

To examine whether the determined affinities reflect binding at the established pockets and to assess the functional relevance of individual residues for nucleotide binding and channel activity, we used site-directed mutagenesis to generate a total of eight alanine mutants: M215A, Y295A, R302A, R358A, the double mutant R302A/R358A, R1433A, Y1485A, and N1487A. These residues were selected based on previous studies and structural analysis (Figure 3). In particular, Huang and coworkers reported that M215A, Y295A, R302A/R358A, and R1433A significantly reduced *hs*TRPM2 currents in response to ADPR. We have recently shown that *hs*TRPM2 R302A/R358A and *hs*TRPM2 R1404Q also do not respond to dADPR^22^. In addition these residues are highly conserved across species, highlighting their likely functional importance^16^. Y1485 is less conserved, with a tyrosine in two of three species and a cysteine in the third, and the mutation Y1485A caused only a mild reduction in current^16^. Although no experimental data were previously available for N1487, structural evidence suggested it could interact with key channel regions, making it a candidate for our study. In the Cryo-EM structure (PDB IDs: 6PUS)^16^, it is located within 3.1 Å of the bound ligand, suggesting that N1487 may be part of the binding site. Structural modeling also indicates a potential hydrogen bond interaction between N1487 and D1431 (Figure 3), which could help stabilize the loop containing Y1485 - residue that forms a π–π interaction with the adenine moiety of the ADPR purine base. However, given the moderate resolution of the structure (3.7 Å), these interactions cannot be assigned with complete certainty.

We first confirmed that all mutants were structurally intact by measuring their melting temperatures (T_m_), which showed no major destabilization compared to the wild-type (WT) protein. Next, we measured binding affinities (K_d_) for ADPR and dADPR using nDSF. Mutagenesis revealed distinct contributions of individual residues to nucleotide binding in TRPM2 (Figure 2). M215A exhibited reduced affinities (K_d_: ADPR = 17 µM [11 µM; 27 µM], dADPR = 2 µM [1.6 µM; 2.9 µM]), indicating that M215 contributes to nucleotidebinding. Y295A caused a marked disruption: while ADPR binding showed a K_d_ of 1.8 µM [0.3 µM; 8.9 µM], dADPR binding was reduced roughly eighty fold with K_d_ = 18 µM [8.6 µM; 37 µM]. The ADPR binding curve was noisy with a wide asymmetric confidence interval, suggesting that Y295 is important for stabilizing ligand interactions.

**Figure 2.**
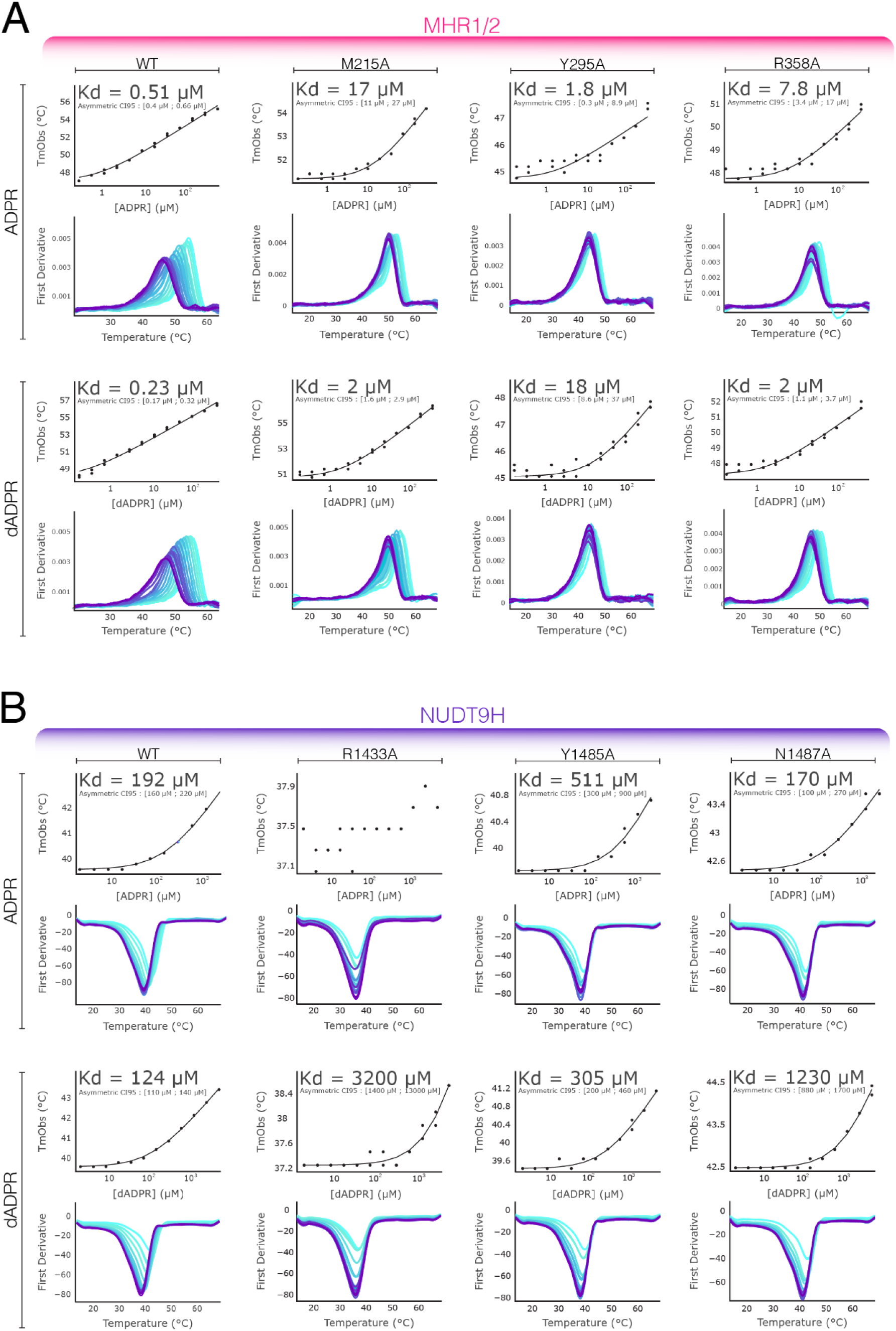
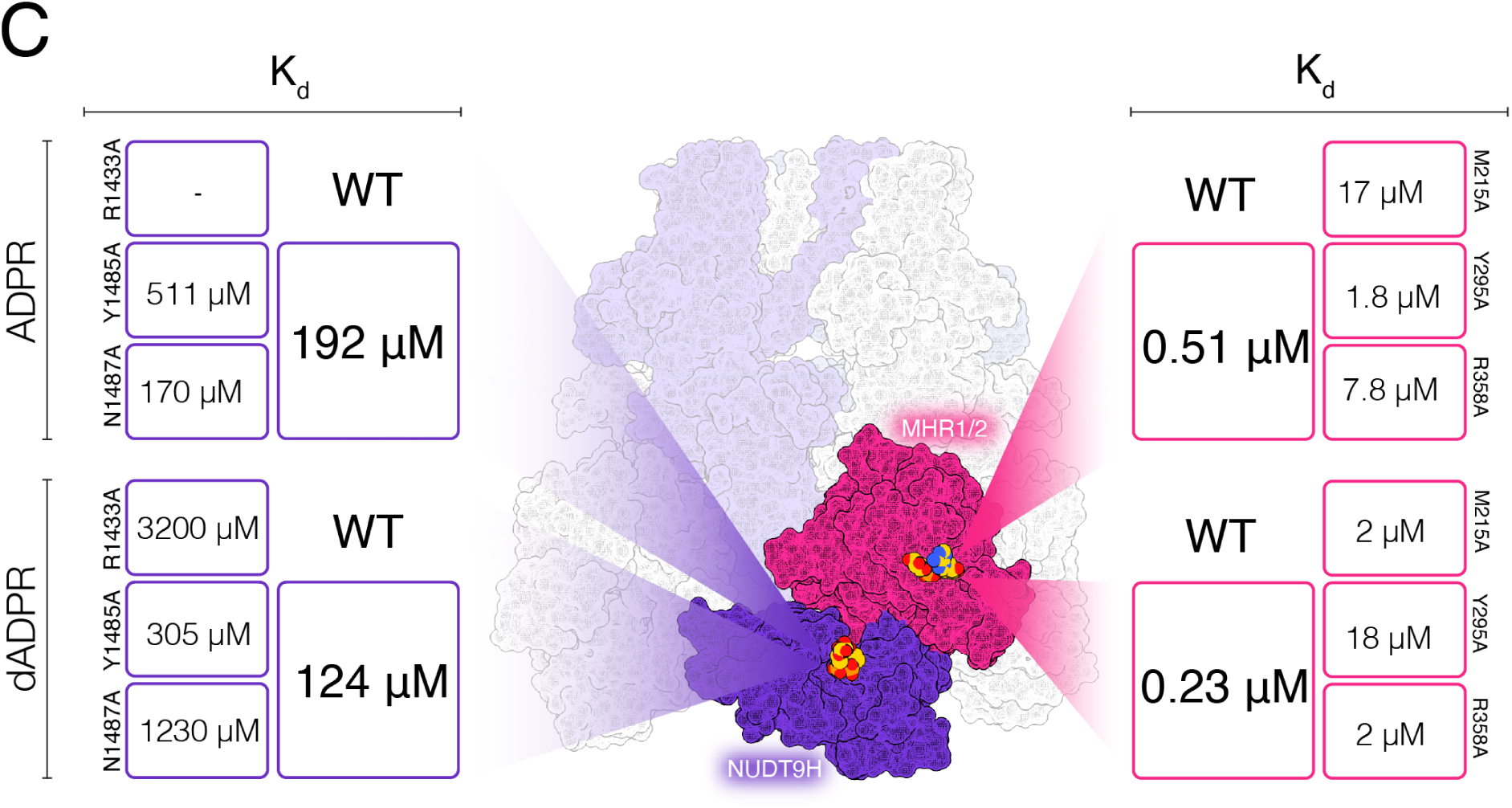
Binding analysis of TRPM2 alanine mutants. Six residues (M215, Y295, R358, R1433, Y1485, N1487) were mutated to alanine to probe their role in ADPR and dADPR binding. nDSF binding assays revealed distinct contributions of individual residues to nucleotide binding in TRPM2. (A) M215A exhibited reduced affinities, indicating that M215 supports nucleotide binding. Y295A caused a marked disruption: the ADPR binding curve is noisy with a wide asymmetric confidence interval, suggesting that Y295 is important for stabilizing ligand interactions. R358A retained partial binding (K_d_: ADPR = 7.8 µM, dADPR = 2 µM), indicating a supportive role in ligand interaction. (B) R1433A nearly abolished binding, highlighting its essential role in the NUDT9H domain. Y1485A and N1487A showed only modest effects. Together, the data confirm that the measured K_d_ values reflect nucleotide binding and identify Y295 and R1433 as key determinants of ligand recognition. In nDSF experiments, where the peak corresponds to the observed melting temperature (T_mObs_), i.e., the temperature at which protein unfolding occurs. T_mObs_ against ligand concentration depicts the ligand-induced stabilization of the protein; a concentration-dependent shift in T_mObs_ reflects ligand binding and is used to calculate the dissociation constant. (C) Graphic representation of comparative K_d_ values for ADPR and dADPR binding to MHR1/2, NUDT9H, and their respective mutants.

R302A and the double mutant R302A/R358A could not be purified, suggesting that R302 may play an important role in maintaining protein structure. R358A retained partial binding (K_d_: ADPR = 7.8 µM [3.4 µM; 17 µM], dADPR = 2 µM [1.1 µM; 3.7 µM]), indicating a supportive role in ligand interaction.

Although the mutations M215A, Y295A and R358A did reduce the affinity for ADPR, the K_d_ was still well below the K_d_ of the NUDT9H domain. Huang et al. had shown previously that M215A and Y295A as well as the double mutant R302A/R358A did abrogate the TRPM2 current evoked by 100 µM ADPR in inside-out patch-clamp experiments^16^. While the effect of the double mutant might have been due to structural destabilization, our results for the M215A and Y295A confirm the importance of the ADPR binding to the MHR1/2 for the activation of *hs*TRPM2.

Mutation of R1433 in the NUDT9H domain to alanine resulted in reduced binding, with an indeterminate K_d_ for ADPR and an affinity reduced by more than twentyfold affinity for dADPR (K_d_ = 3200 µM [1400 µM; 13000 µM]), highlighting its essential role in the NUDT9H domain.

Y1485A exhibited only modest increases in K_d_ (ADPR = 511 µM [300 µM; 900 µM], dADPR = 305 µM [200 µM; 460 µM]), suggesting a minor contribution to binding, consistent with lower evolutionary conservation. N1487A showed no major change for ADPR (K_d_ = 170 µM [100 µM; 270 µM]) but a tenfold reduction in affinity for dADPR (K_d_ = 1230 µM [880 µM; 1700 µM]), indicating it is less critical than R1433 but plays a role in ligand recognition. Although residue N1487 is not positioned directly within the 2′-deoxy binding site, its mutation has a stronger effect on dADPR binding than on ADPR binding. One possible explanation is that the 2′-OH group of ADPR can form a hydrogen bond with the OH-group of a Y1485, which may help stabilize the adjacent helix. This stabilizing interaction is absent with dADPR, potentially increasing the sensitivity of the loop to further destabilizing mutations, such as N1487A. Notably, whole-cell patch-clamp recordings showed that N1487A did not significantly affect activation by either ADPR or dADPR (Figure 3). This might be due to the difference in measurement temperature between the assays. Binding affinity (K_d_) was determined at higher temperature (∼ 43 °C), whereas patch-clamp recordings were performed at 25 °C. We speculate that at higher temperatures, the helix containing residue N1487 may be more dynamic. In this context, loss of the hydrogen between Y1485 and dADPR bond together with the N1487A mutation could further enhance local flexibility and destabilize the interaction. At 25 °C, reduced molecular motion may partially compensate for these effects, resulting in minimal functional differences in the electrophysiological measurements.

**Figure 3.**
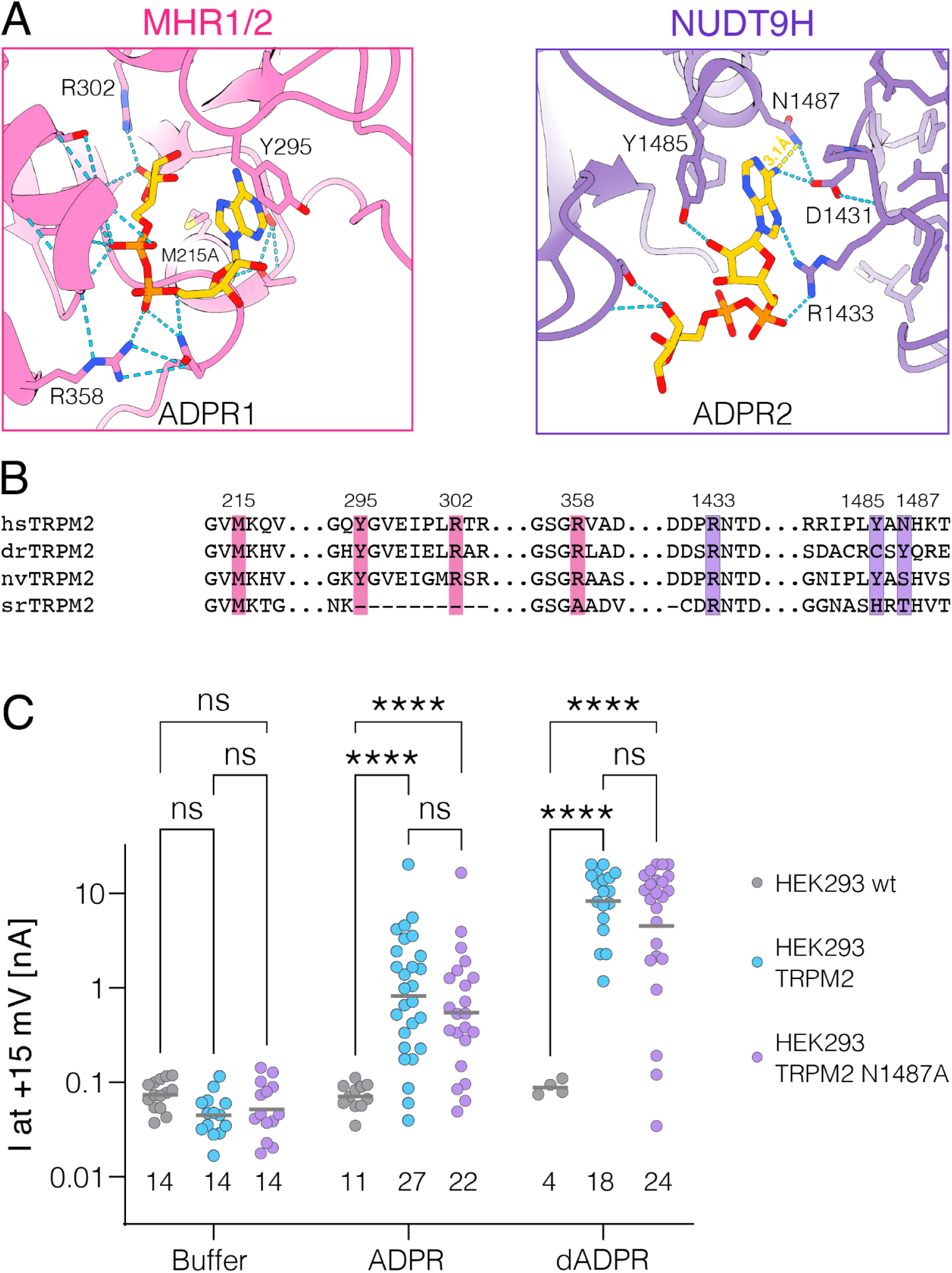
The TRPM2 mutant N1487A exhibits no significant effect on ADPR or dADPR-induced currents compared to wild-type TRPM2. (A) Cryo-EM structure of the TRPM2 MHR1/2 and NUDT9H domains in complex with ADPR (PDB ID: 6PUS)^16^. The protein is shown in cartoon representation, with ADPR rendered as sticks. Key amino acid residues analyzed in this study are highlighted. Hydrogen bonds identified in the deposited structure are shown as cyan dashed lines. Distances between ADPR and N1487 are shown as yellow dashed lines and were measured using ChimeraX (3.1 Å). (B) Sequence alignment of TRPM2 orthologs from human (*Homo sapiens* TRPM2), zebrafish (*Danio rerio* TRPM2), the sea anemone (*Nematostella vectensis* TRPM2), and the choanoflagellate (*Salpingoeca rosetta* TRPM2). Residues analyzed in this study are highlighted. (C) HEK293 cells were transiently transfected with either wild-type TRPM2 (pink) or TRPM2 mutant N1487A (purple), while non-transfected HEK293 cells served as a negative control (gray). Whole-cell patch-clamp recordings were performed 24 h post-transfection. The intracellular pipette solution contained either 100 µM ADPR, dADPR, or no nucleotide (buffer). Each data point represents the maximum whole-cell current at +15 mV from an individual cell. Numbers below the scatter plot indicate the total number of cells recorded per condition. Gray bars denote the mean of the log-transformed data. Log-transformed data were analyzed using two-way ANOVA and Tukey’s multiple comparisons test (****p < 0.0001; ns, not significant).

Overall, these results confirm that the K_d_ values determined reflect binding of ADPR to the established sites and show that specific residues, such as Y295 and R1433, are essential for interaction with the nucleotide. Interestingly, some mutations affected ADPR and dADPR binding differentially, suggesting that the two nucleotides may engage the binding sites in distinct ways.

### Cellular ADPR levels suggest agonist binding to MHR1/2 triggers activation of TRPM2

The large difference in affinity between the two nucleotide-binding sites—spanning approximately three orders of magnitude—raises important questions regarding the physiological concentration of ADPR in resting versus stimulated cells. If basal ADPR levels are already sufficient to saturate the high-affinity MHR1/2 domain, ADPR may play a structural or stabilizing role at this site. This could explain the pronounced effects of point mutations within MHR1/2, while suggesting that this binding event alone does not directly drive hsTRPM2 activation under physiological conditions. In such a scenario, channel activation would require an increase in ADPR concentration to levels that permit binding to the lower-affinity NUDT9H domain. Conversely, if the resting ADPR concentration is below the Kd of the MHR1/2 site, stimulation-induced increases in ADPR would first promote binding to the high-affinity MHR1/2 domain. Only upon further elevation of intracellular ADPR levels would occupancy of the low-affinity NUDT9H site occur, potentially triggering channel activation.

We determined the basal ADPR concentration in HEK293T cells using metabolomics, yielding a value of 0.39 µM (Supplementary Fig. 5). After 3 minutes of incubation with H₂O₂, the measured ADPR concentration decreased to 0.16 µM, and no increase in ADPR levels was observed. This may reflect transient changes occurring within a narrow time window that are not captured under our experimental conditions.

As metabolomics (and earlier methods such as 2D-HPLC) require nucleotide extraction from intact cells, we complemented these measurements with an orthogonal approach using a fluorescent ADPR biosensor^27^ (Fig. 4). This biosensor is based on NrtR, an ADPR-responsive transcriptional repressor from soil bacterium *Shewanella onedensis*^21^. Circularly permutated Venus (cpVenus) was inserted into an engineered variant of NrtR at an allosteric position responsive to ligand binding^28^. The biosensor was co-expressed as a fusion protein with mCardinal to enable a ratiometric readout..

**Figure 4.**
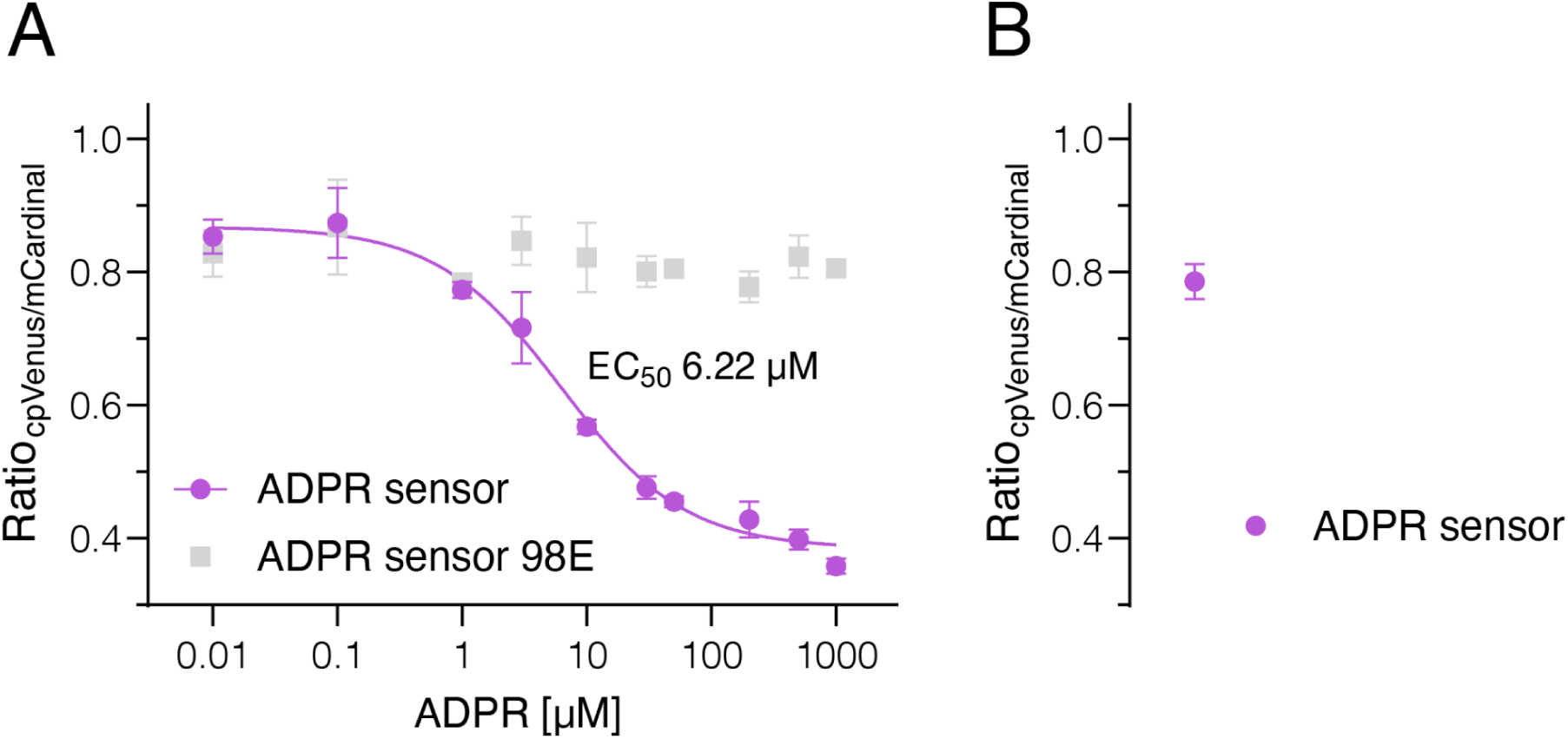
ADPR concentration-response curve of the ADPR biosensor and an ADPR-insensitive mutant 98E. (A) ADPR was added at different concentrations to permeabilized HEK293T cells, either lentivirally transduced with the ADPR biosensor (purple) or the mutant 98E (grey). Fluorescence was measured at 475/515 nm and 604/659 nm (excitation/emission) at 37 °C. (B) Intact cells were used to determine basal ADPR levels. Measurements were performed in a 96-well plate with three technical replicates per condition and four areas per well. Mean values were calculated from three to five independent experiments. Data were analyzed using nonlinear regression ([agonist] vs. response – variable slope, four parameters), resulting in an EC₅₀ of 6.22 µM for the ADPR biosensor (minimum asymptote: 0.3849; maximum asymptote: 0.8684; Hill slope: –0.8719). Error bars represent the SEM.

To determine intracellular ADPR concentrations, we performed an in situ calibration in saponin-permeabilised cells using defined ADPR concentrations. A four-parameter logistic function was fitted to the resulting data, and, based on the parameters of the concentration–response curve (Fig. 4), we estimated the basal free ADPR concentration in intact cells to be approximately 1.0 µM.

In the past hydrogen peroxide has been used consistently to activate TRPM2 in HEK293 and 293T cells via the PARP/PARG pathway^29–34^, suggesting that the resulting ADPR concentrations are sufficient for TRPM2 activation. To determine the intracellular ADPR concentrations reached during this process, we exposed HEK293T cells transduced to express the ADPR biosensor to various concentrations of hydrogen peroxide and monitored ADPR production over a period of 20 minutes (Fig. 5A). Recognisable ADPR production began at 50 µM H₂O₂ and plateaued between 500 µM and 1 mM (Fig. 5B). Calibration of the recordings showed that the total cellular ADPR concentration increased from approximately 1 µM to 10 µM before decreasing again. Pre-incubation with PJ34 abolished the signal, confirming the dependence of ADPR formation on PARP activity.

**Figure 5.**
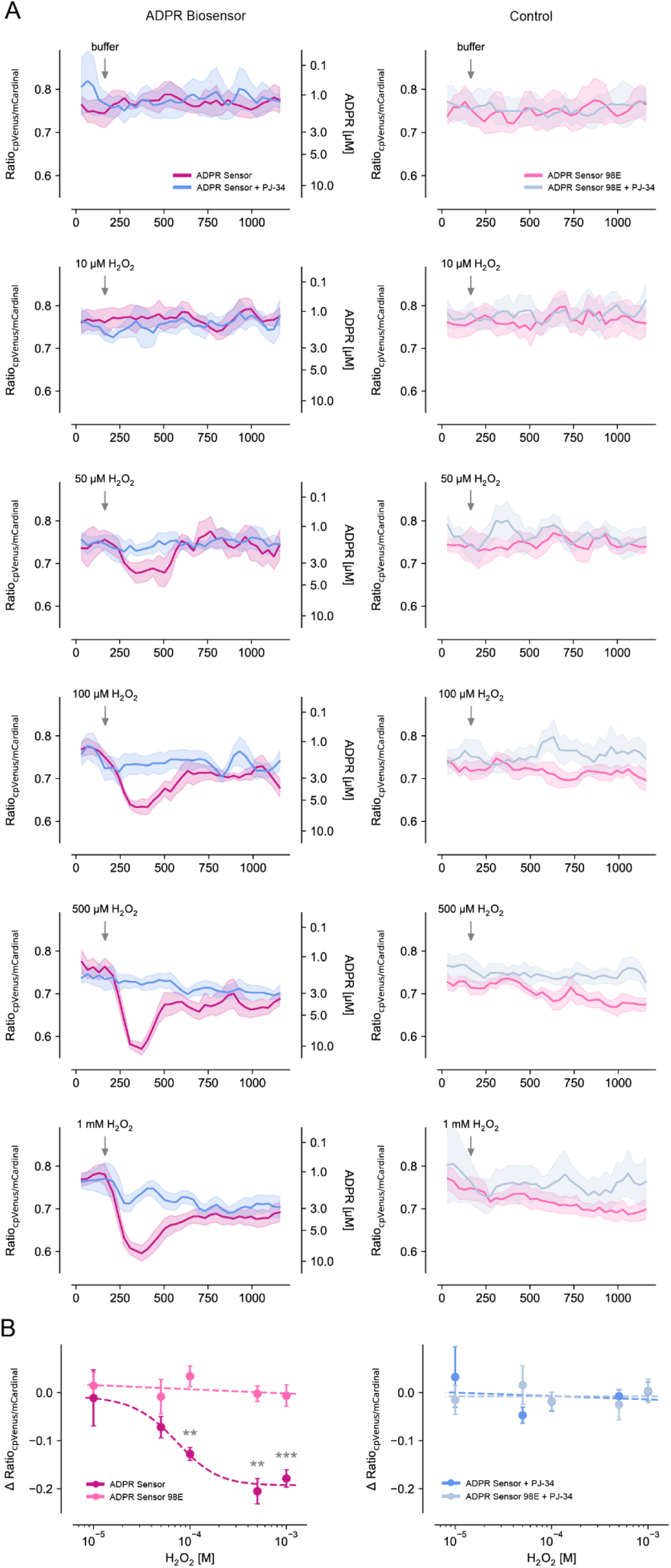
Time course of intracellular ADPR after exposure to hydrogen peroxide determined using the ADPR biosensor. A) HEK293T cells were lentivirally transduced with either the ADPR biosensor (left) or the ADPR-insensitive mutant 98E (right) and treated with buffer or varying concentrations of H₂O₂ at 37 °C at the indicated time point. Measurements in the presence of the PARP inhibitor PJ34 (10 µM) are shown in blue. Fluorescence was recorded at 475/515 nm and 604/659 nm (excitation/emission). Experiments were performed in 96-well plates with two technical replicates per condition and four regions per well. Mean values were calculated from four independent experiments and a Savitzky-Golay filter (window of 3) was applied. The ADPR concentration on the right axis of the left panels was calculated using the minimum (0.3849) and maximum (0.8684) asymptotes, as well as the Hill slope (–0.8719) obtained from the ADPR concentration–response curve of the biosensor (Figure 4). Data are presented as mean ± SEM. B) Concentration dependent change in Ratio after exposure to H_2_O_2_. ΔRatio was calculated from the average before (100 to 150 s) and after exposure (350 to 400 s) to H_2_O_2_ and tested between sensor and mutant using Welch’s t-test. **p < 0.01; *** p< 0.001; n.s., not significant.

Both, the concentration in resting cells is lower than what has been previously determined for Jurkat cells (44 µM) and murine BW5147 cells (73 µM) using 2-dimensional HPLC^35^. This difference might either reflect cell type specific differences, e.g. in the expression of CD38 or other metabolic enzymes, a potential hydrolysis of poly-ADPR or NAD during cell lysis and nucleotide extraction in the older study, or release of ADPR bound to proteins or cellular structures during the extraction procedure.

## Discussion

Previous studies have reported a wide range of apparent ADPR sensitivities for TRPM2, with EC₅₀ values spanning submicromolar to hundreds of micromolar. For recombinant TRPM2 expressed in HEK293 cells, whole-cell patch-clamp recordings performed at room temperature show that channel activation occurs in the micromolar range. In the absence of cytosolic Ca²⁺, activation requires relatively high agonist concentrations (approximately 40–90 µM)^12,36,37^. In contrast, when cytosolic Ca²⁺ is present and unbuffered, activation is observed at much lower concentrations, around 12 µM^21^. It is important to note that HEK293 cells do not naturally express TRPM2. In immune cells, activation starts to appear at even lower ADPR concentrations under unbuffered Ca²⁺ conditions, with activation detected at ∼10 µM for Jurkat cells and 0.3–1 µM for neutrophils^38–40^. By comparison, in monocytes, activation requires much higher ADPR concentrations (∼130 µM) in the absence of Ca²⁺^12^. Notably, reported intracellular ADPR levels — ∼40 µM in Jurkat cells and 5 µM in neutrophils — exceed these minimal activation thresholds^10,35,39^. These observations suggest that TRPM2 would be expected to exhibit constitutive activity in these cells, unless additional mechanisms constrain channel activation.

Our measured K_d_ values for the separately expressed domains, MHR1/2 = 0.5 µM and NUDT9H = 192 µM, provide a possible explanation for this behavior. The high-affinity MHR1/2 sensor can be occupied at sub- to low-micromolar ADPR concentrations, potentially priming the channel or inducing slow opening. In contrast, the much lower-affinity NUDT9H site would require tens to hundreds of micromolar ADPR to become substantially occupied. However, such concentrations are unlikely to be achieved under physiological conditions in vertebrate cells. Even during oxidative stress induced by H₂O₂, the total cellular NAD pool is typically limited to the low hundreds of micromolar range^41–43^, and only a fraction of this pool can be converted to ADPR. In addition, ADPR generated in the nucleus must diffuse through the cytosol to reach the plasma membrane, where it is likely subject to degradation by NUDT5 and potentially cytosolic NUDT9 isoforms. These factors are expected to strongly constrain the accumulation of high ADPR concentrations at the membrane.

Taken together, these considerations suggest that, in vertebrate TRPM2 channels, the NUDT9H domain is unlikely to function as a physiologically relevant ligand-binding site. Instead, its role may be primarily structural, reflecting its tight integration into the channel architecture, while channel regulation under physiological conditions is predominantly mediated by high-affinity binding to the MHR1/2 domain. Several observations support this view. First, the affinity of the isolated NUDT9H domain for ADPR is low (K_d_ ≈ 192 µM), whereas total cellular ADPR concentrations following stimulation with 1 mM H₂O₂, sufficient for full channel activation, do not exceed ∼10 µM. In HEK293 cells, ADPR is generated predominantly in the nucleus via PARP/PARG activity, and membrane-associated CD38 does not contribute to this pathway. Thus, local concentrations at the plasma membrane substantially exceeding total cellular ADPR levels appear unlikely. Unless tetrameric channel assembly dramatically increases the affinity of the NUDT9H domain, this site would not be expected to be significantly occupied at cellular ADPR concentrations well below ∼40 µM (i.e., <1/5 of the measured K_d_).

Second, the presence of ADPR in the NUDT9H domain in cryo-EM structures does not necessarily imply physiological occupancy, as these structures were obtained in the presence of 1 mM ADPR—conditions sufficient to force binding to a low-affinity site.

Third, functional measurements are not easily reconciled with a primary gating role of NUDT9H binding. Reported EC_50_ values for ADPR activation typically range from 1 to 90 µM^44^ (we determined an EC_50_ of 28 µM)^10^, which is inconsistent with a K_d_ of ∼192 µM. The discrepancy between the affinity of the (isolated) MHR1/2 domain and the EC_50_ of the tetrameric channel is reminiscent of the relationship between myoglobin and hemoglobin: while a single MHR1/2 domain binds agonists tightly, the tetrameric TRPM2 channel displays cooperative behaviour (n_H_ ≈ 5.5 with unbuffered Ca²⁺)^45^, so that multiple binding events are required to open the channel. As a result, the macroscopic current response occurs at higher agonist concentrations than the intrinsic affinity of a single domain would suggest.

Fourth, although mutations within the NUDT9H domain (e.g., R1433A) have been interpreted as evidence that ADPR binding at this site is required for activation, several findings argue for a structural interpretation. We have shown that W1355A in NUDT9H alters activation by dADPR but only slightly affects ADPR responses, despite being located outside the canonical binding pocket^11^. Ehrlich et al. report that N1326D, also outside the binding pocket, disrupts interdomain interactions and prevents channel activation^19^. Moreover, mutations that significantly reduce ligand affinity at NUDT9H (e.g., Y1485A: 192 µM → 511 µM; N1487A: 124 µM → 1230 µM for dADPR) have only minor or no effects on channel activation at 25 °C. These observations are difficult to reconcile with a model in which ligand binding to NUDT9H is required for gating. Notably, NUDT9H is less thermally stable, as its melting temperature (T7 = 40 °C) is lower than that of MHR1/2 (T7 = 49 °C). This lower stability suggests that, although NUDT9H may not directly bind ADPR to trigger gating, it could interact with CaM as part of temperature sensing, consistent with our finding that NUDT9H must be thermally destabilized for this interaction^46^.

The most consistent interpretation is that binding of ADPR or dADPR to the MHR1/2 domain is sufficient to induce channel gating, whereas the NUDT9H domain, although required for structural integrity and likely contributing to conformational transitions during gating, is not substantially occupied under physiological conditions. A strictly sequential binding model is difficult to reconcile with the high ligand concentrations required to engage the NUDT9H site, unless additional assumptions are invoked, for example, that ligand binding to MHR1/2 markedly increases the affinity of NUDT9H. This interpretation is further supported by our quantitative framework, which shows that MHR1/2 is expected to be occupied within the physiological to oxidative stress range of ADPR concentrations, whereas NUDT9H remains largely unoccupied due to its substantially lower affinity (Fig. 6A).

**Figure 6.**
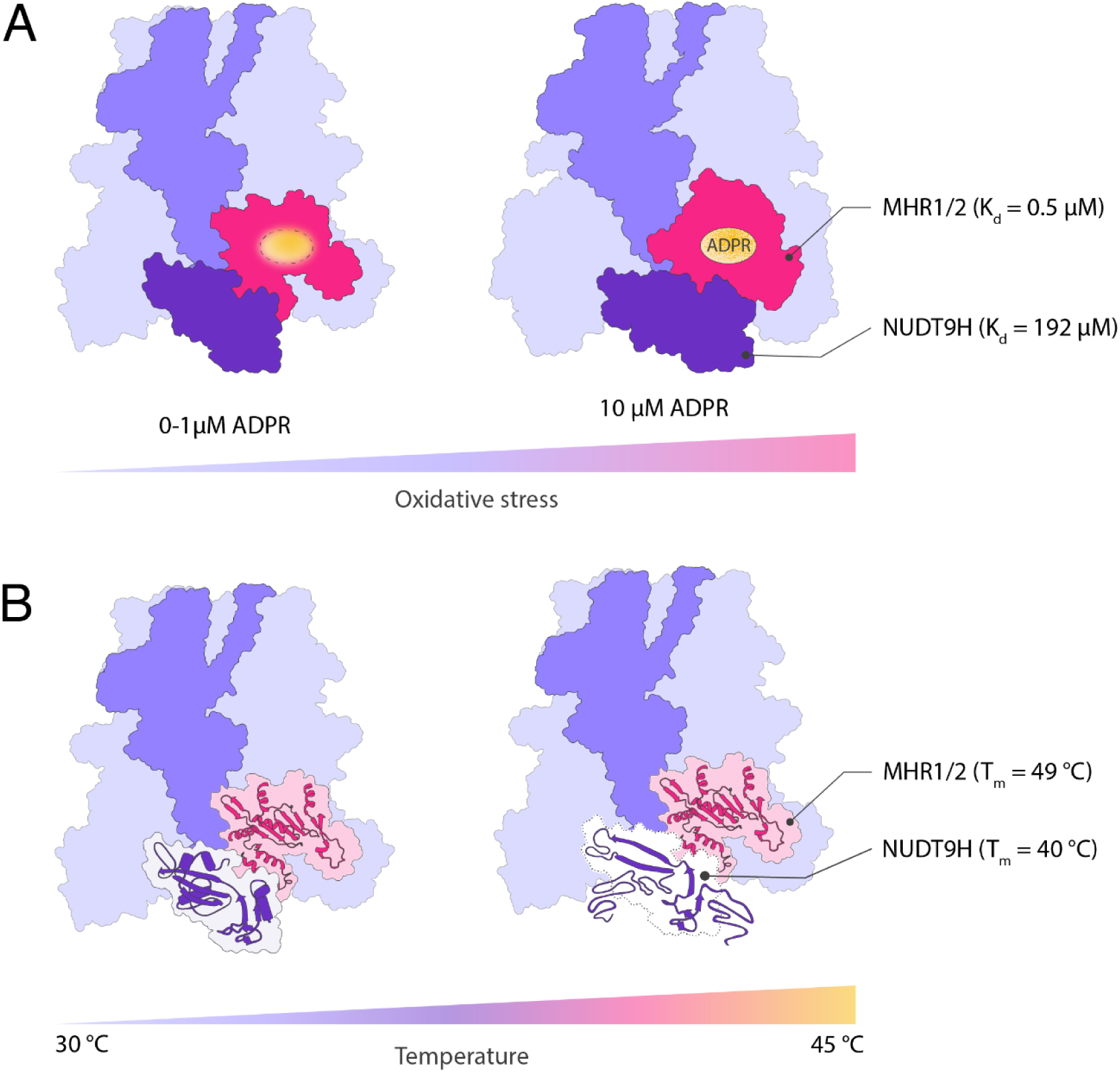
Temperature and oxidative stress effects on TRPM2. **(A)** Under physiological conditions, the ADPR concentration near TRPM2 is approximately 1 µM. With increasing oxidative stress, ADPR levels can rise to around 10 µM. Based on the dissociation constants (K_d_), MHR1/2 may be partially occupied under physiological conditions but is expected to be fully occupied under oxidative stress (K_d_ = 0.5 µM). In contrast, the NUDT9H binding site is unlikely to bind ADPR under either condition (K_d_ = 192 µM). (B) Thermal stability analysis shows that NUDT9H has a lower melting temperature (T_m_ = 40 °C) compared with MHR1/2 (T_m_ = 49 °C), indicating that NUDT9H is less thermostable and may partially unfold under elevated temperatures.

It should be noted, however, that the intracellular spatial distribution of ADPR remains incompletely understood. Localized ADPR production could create microdomains with higher concentrations than bulk measurements suggest. CD38, a membrane-associated enzyme, may generate elevated local ADPR concentrations and could contribute to TRPM2 regulation in immune cells during inflammatory responses, even though this mechanism does not explain H₂O₂-induced activation in HEK293 cells. In addition, the lower thermal stability of the NUDT9H domain compared with MHR1/2 (Fig. 6B) suggests that it may be more susceptible to conformational flexibility or partial unfolding under stress conditions, potentially modulating its structural role during channel activation.

## Materials and Methods

### *hs*MHR1/2 expression and purification

The coding sequence for residues 53–446 of human TRPM2 (*hs*TRPM2) was cloned into the pXLG-cterm-GFP-His vector using SLiCE cloning at the EcoRV restriction site. A TEV protease cleavage site was inserted between *hs*MHR1/2 and the C-terminal GFP-His tag to facilitate downstream purification. Expi293 cells (suspension-adapted HEK293 cells; Thermo Fisher) were cultured in FreeStyle 293 Expression Medium at 37 °C with 8% CO₂, shaking at 220 rpm with a 1-inch orbital diameter. Cells were maintained at a density of 0.3–4 × 10⁶ cells/mL by passaging every 2–3 days. Cells were transiently transfected with pXLG-cterm-GFP-His expression vectors containing 53–446 of *hs*TRPM2 variants (wild type, M215A, Y295A, R302A, R358A and R302/358A double mutant). For transfection, cells were concentrated to 20 × 10⁶ cells/mL and transfected using linear 40 kDa polyethylenimine (PEImax) at a ratio of 1 µg plasmid DNA and 3 µg PEImax per 10⁶ cells. The transfection mixture was incubated for 1 hour with shaking, after which fresh medium was added to adjust the cell density to 1 × 10⁶ cells/mL. Cells were incubated for 24 hours at 37 °C, followed by the addition of sodium butyrate to a final concentration of 10 mM, and cultured for an additional 48 hours at 30 °C.

The cells were resuspended in 5mL/g pellet buffer L (50 mM NaP pH 7.3, 290 mM NaCl, 10 mM KCl, 10% glycerol v/v, 0.5 mM Tris(2-carboxyethyl)phosphine (TCEP), 10 mM imidazole and EDTA-free protease inhibitor (Roche, Germany)) and lysed with a high-pressure homogenizer (EmulsiFlex-C3, Avestin). The lysate was cleared by centrifugation (18,000 x *g*, 30 min) and applied to Ni^2+^-resin (Ni-NTA Agarose, QIAGEN). After elution with 250 mM imidazole, removal of the GFP-His tag with TEV-protease and dialysis overnight a reverse IMAC was performed and the protein then further purified by gel filtration using a Superdex 200 Increase 10/300 GL with buffer SEC (50 mM NaP pH 7.3, 290 mM NaCl, 10 mM KCl, 0.5 mM TCEP).

### *hs*NUDT9H expression and purification

The sequence coding for the residues 1236-1503 of *hs*TRPM2 was cloned into a pQE-80 vector with a N-terminal 6xHis-tag using the *Sph*I and *Hind*III restriction sites. The expression was carried out by growing transformed BL21 gold pLysS *E.coli* to an OD of 0.5 at 37 °C in LB, inducing with 0.4 mM isopropyl β-D-1-thiogalactopyranoside (IPTG), lowering the temperature to 30 °C and harvesting after 5 hours by centrifugation (24 min, 4,000 x *g*). The pellet was resuspended in buffer L (50 mM Tris-HCl pH 7.5, 300 mM NaCl, 0.5 mM TCEP, EDTA-free protease inhibitor and lysed with a high-pressure homogenizer (EmulsiFlex-C3, Avestin). The lysate was cleared by centrifugation (18,000 x *g*, 60 min) and applied to Ni^2+^-resin. After elution with 250 mM imidazole the pooled fractions were dialyzed overnight into buffer SEC (50 mM Tris-HCl pH 7.5, 0.5 mM TCEP) and applied to a HiTrap SP HP (Cytiva) column. Elution was performed as a gradient with buffer B(cx)(50 mM Tris-HCl pH 7.5, 1 M NaCl, 0.5 mM TCEP). The purification was finalized by gel filtration on a Superdex200 Increase 10/300 GL with buffer SEC (50 mM Tris-HCl pH 7.5, 0.5 mM TCEP).

### Differential scanning fluorimetry (nDSF)

Thermal stability measurements were performed using a Prometheus NT.48 instrument (NanoTemper Technologies) to assess ligand binding. A 12-point, 2-fold serial dilution of ADPR or dADPR was prepared, ranging from 0.2 µM to 400 µM. Each ligand concentration was mixed with *hs*MHR1/2 protein at a final concentration of 1 µM in buffer N (50 mM Tris-HCl, pH 7.5, 0.5 mM TCEP). Protein unfolding was monitored by tracking changes in intrinsic tryptophan fluorescence during a thermal ramp from 15 °C to 70 °C at a rate of 1 °C/min. For a qualitative assessment of ligand binding to the *hs*NUDT9H domain, 5 mM of ligand was added to 5 µM protein. Ligand binding was assessed using the unfolded fraction fit in FoldAffinity^47^.

### Isothermal titration calorimetry (ITC)

ITC measurements were carried out at 25°C using a MicroCal PEAQ-ITC isothermal titration calorimeter (Malvern Panalytical) and thermodynamic parameters were analyzed using the MicroCal ORIGIN™ software. The ligands (ADPR, dADPR, 8-Br-ADPR) were each added from an aqueous stock (100 mM for ADPR, 100 mM dADPR, 50mM for 8-Br-ADPR) for to the corresponding SEC buffer (see purification) to a concentration of 2 mM for *hs*NUDT9H and placed in the syringe. After an initial injection of 0.4 µL, 18 regular injections of 100 µM *hs*NUDT9H in the sample cell. The individual injections were interspaced by 150 s and stirring speed was set to 750 rpm. Heat of dilution was obtained by titrating the ligands into the corresponding SEC buffers and baseline corrections were carried out accordingly. All ITC experiments were performed as triplicates and errors are reported as standard deviations of the mean K_d_ value.

### Cell culture

The HEK293 cells used for patch-clamp experiments were obtained from the DSMZ (ACC305, Braunschweig, Germany). HEK293T cells lentivirally transduced with the ADPR biosensor were provided by Xiaolu Lim Ang Cambronne (UT Austin, USA). All cells were cultured in DMEM with 4.5 g/L D-glucose and Glutamax-I (Thermo Fisher Scientific, Germany), supplemented with 10% FBS (Capricorn Scientific, Ebersdorfergrund, Germany), 100 units/mL penicillin, and 100 µg/mL streptomycin (Thermo Fisher Scientific, Germany). Additionally, 2 µg/mL of the selection antibiotic puromycin (Thermo Fisher Scientific Gibco, Germany) was added to the medium of the ADPR biosensor and mutant 98E cells. Cell cultures were maintained at 37 °C in a humidified atmosphere and 5% CO_2_.

### Generation of stable HEK293T cell lines with ADPR biosensor

HEK293T cells (500,000 cells per well) were seeded in 6-well TC-treated plates (Fisherbrand) and cultured in Dulbecco’s modified Eagle’s medium (DMEM, Sigma-Aldrich) containing 10% Fetal bovine serum (FBS, Biowest) at 37 °C with 5% CO_2_. Cells were transfected at ∼90% confluency the next day using polyethylenimine (PEI) at a ratio of 5:1 PEI:DNA. The transfection mixture was prepared in Opti-MEM (Gibco) and comprised of 20 μg PEI and a plasmid mixture of 1 μg psPAX2, 1 μg pMD2.G, and 2 μg of either pLenti-NES-Flag-HA-ADPR Sensor-mCardinal-IRES-Puro or the pLenti-NES-Flag-HA-R98E-mCardinal-IRES-Puro control plasmids. Supernatant that contained viral particles was collected 72 hours post-transfection and filtered through a 0.45 μm PVDF filter (Millex). Five hundred microliters of viral supernatant were incubated with a cell suspension of HEK293T cells (125,000 cells/mL) in a 6-well plate for 2 days. Transduced cells were selected with puromycin (2 μg/mL) and cultured with selection antibiotic for 2 days. Approximately 30% of cells exhibited puromycin resistance, suggesting an MOI < 1. Transduced cells were maintained in selection media until all non-transduced control cells succumbed to puromycin selection in parallel. Stable expression was verified with Western Blotting and live cell imaging.

### Mutagenesis

Mutagenesis was performed as described recently^11^. Briefly, QuikChange mutagenesis (QuickChange Site-Directed Mutagenesis Kit, Agilent Technologies) was used to introduce the N1487A mutation into the plasmid pIRES2-TRPM2-EGFP^2^ using the primers 5′-CATCCCACTCTATGCGGCCCACAAGACCCTC-3′ and

5′-GAGGGTCTTGTGGGCCGCATAGAGTGGGATG-3′. The presence of the mutation in the resulting plasmid was confirmed by commercial DNA sequencing (Microsynth, Göttingen, Germany).

For nDSF, mutagenesis was performed via Quick Change PCR. Mutations were introduced into plasmids pXLG_MHR1/2_eGFP_His6 and pQE-80L_His6_NUDT9H using forward (F) and reverse (R) primers indicated in table 1:

**Table 1.**
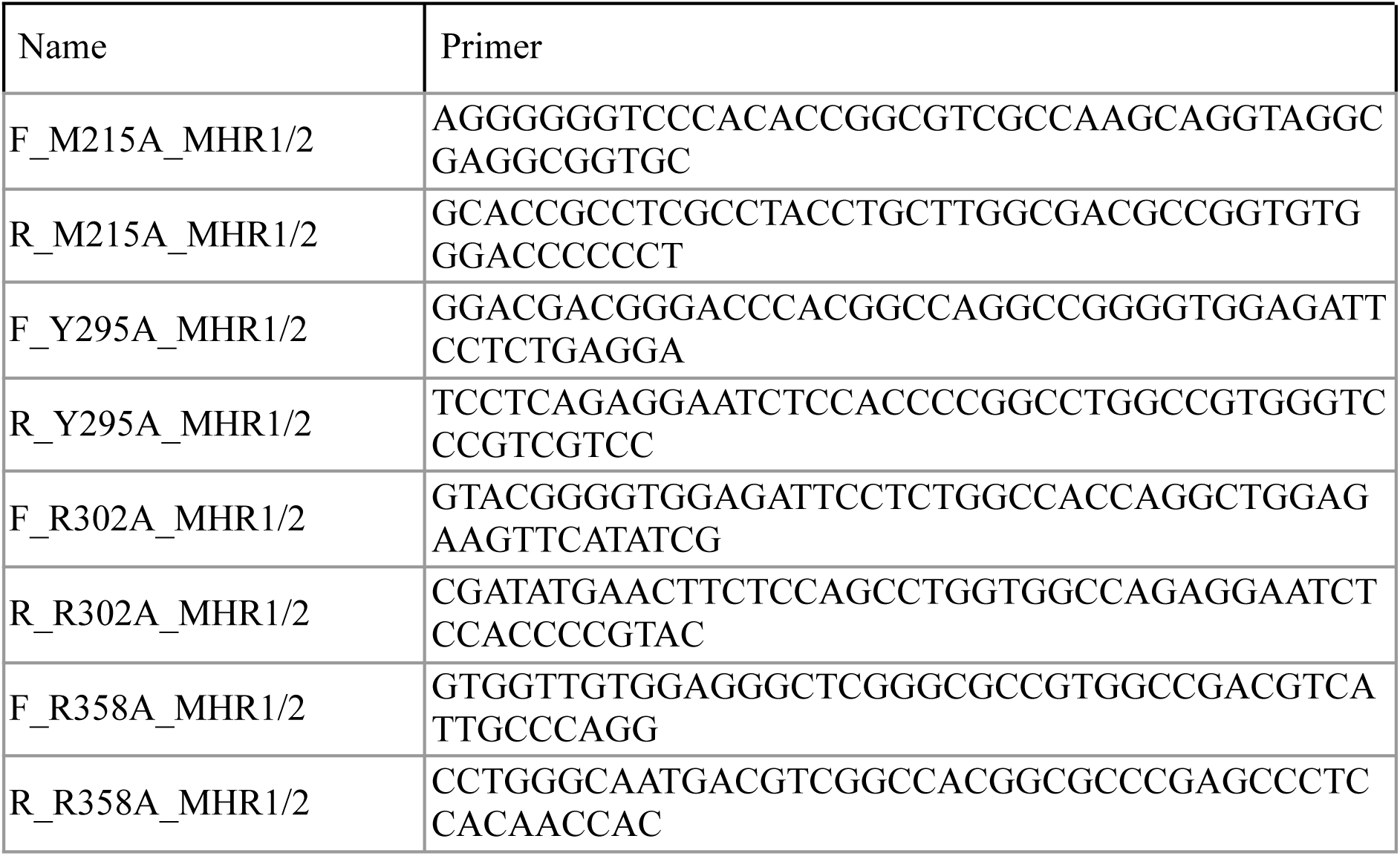
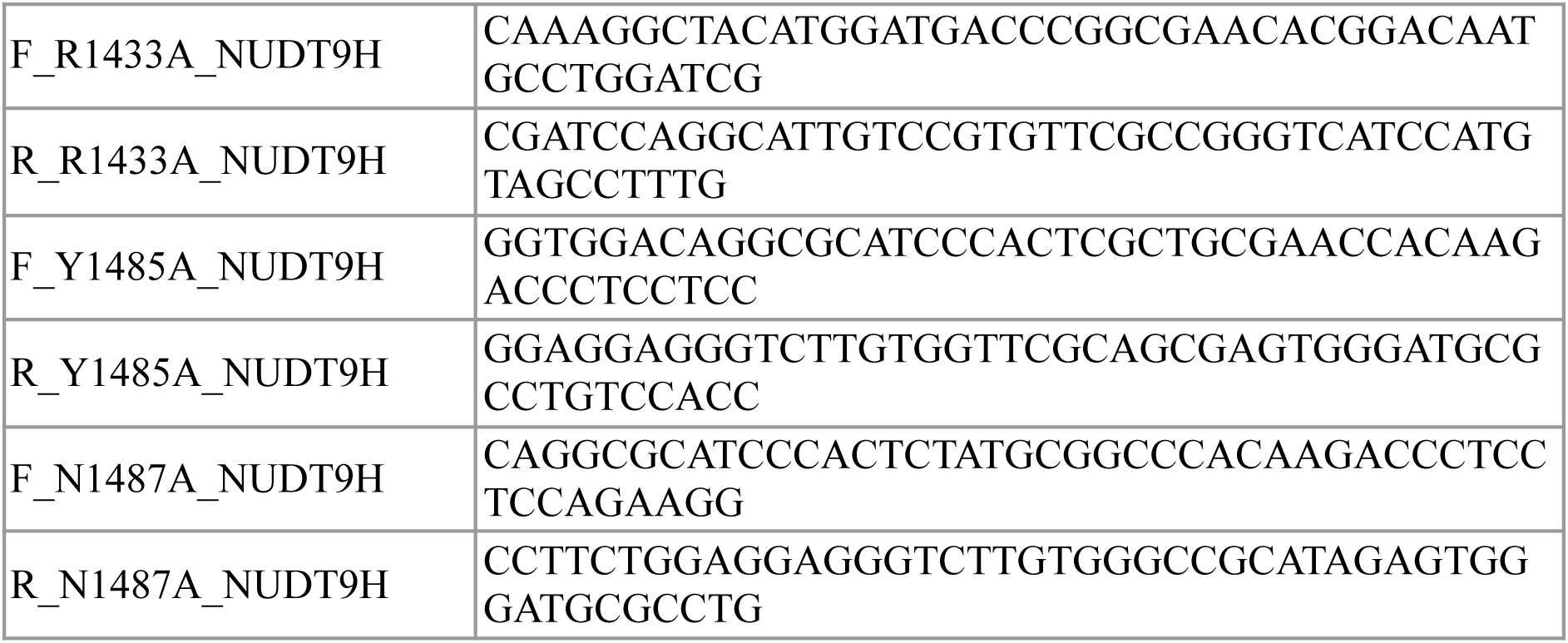
Primers for Quick Change PCR.

### Electrophysiology

Electrophysiology experiments were conducted as described previously^11,22^.In summary, HEK293 cells were transfected in suspension with pIRES2-TRPM2-EGFP^48^ or pIRES2-TRPM2-EGFP N1487A using JetPEI (Sartorius, Göttingen, Germany) and seeded into 35 mm dishes (Greiner Bio-one, Germany) at a cell density of 2.5 x 10^5^ one day prior to the experiments. Immediately before recording, the culture medium was replaced with bath solution containing: 140 mM N-methyl-D-glucamine, 5 mM KCl, 3.3 mM MgCl_2_, 1 mM CaCl_2_, 5 mM glucose, and 10 mM HEPES, adjusted to pH 7.4 with HCl at room temperature. During patch-clamp recordings, cells were maintained at room temperature. Patch pipettes were pulled from borosilicate capillaries (1.05 x 1.50 x 80 mm; Science Products, Germany) using a P-97 horizontal puller (Sutter Instruments, USA). Resulting pipettes had series resistances between 1.0 MΩ and 3.5 MΩ and were filled with intracellular solution containing: 120 mM KCl, 10 mM HEPES, 10 mM EGTA, 8 mM NaCl, 1 mM MgCl_2_, and 5.6 mM CaCl_2_, adjusted to pH 7.2 with KOH at room temperature, resulting in a free Ca^2+^ concentration of 200 nM). Intracellular solution was supplemented with either 100 µM ADPR, 100 µM dADPR, or no nucleotide. A silver chloride pellet (Science Products, Germany) was used as bath electrode. Whole-cell patch-clamp recordings were performed using an EPC-10 USB amplifier (HEKA, Germany) controlled using PatchMaster software (HEKA, v2x92). To monitor TRPM2 activation following break-in, voltage ramps from -85 mV to +20 mV over 140 ms were applied every 5 s, starting from a holding potential of -50 mV for a total duration of 450 s. Series resistance was compensated automatically using PatchMaster software. Data were sampled at 5 kHz and recorded to hard disk. The maximum outward current amplitude from the voltage ramps (at +15 mV) was used as a measure of TRPM2 activity. Cells in which the seal ruptured during recording, had a series resistance >10 MΩ, or showed significant changes in series resistance during the recording were excluded from further analysis. Due to a right-skewed distribution of the current amplitudes, values were log-transformed to approximate normality for subsequent statistical analysis.

### Fluorescence-based measurements of ADPR using an ADPR biosensor

Cells were seeded at a density of 5 × 10⁴ cells per well into 96-well black microplates with glass bottoms (Corning Incorporated, USA). On the day after seeding, the culture medium was replaced with Ca²⁺ measurement buffer containing: 140 mM NaCl, 5 mM KCl, 1 mM CaCl₂, 1 mM MgSO₄, 1 mM NaH₂PO₄, 20 mM HEPES, and 5.5 mM glucose, adjusted to pH 7.55 with NaOH at room temperature to obtain pH 7.4 at 37 °C. Fluorescence measurements were performed at 37 °C using an Infinite 200 Pro fluorescence microplate reader (Tecan Group Ltd., Männedorf, Switzerland). CpVenus and mCardinal fluorescence was recorded at 475/515 nm and 604/659 nm (excitation/emission), respectively, with excitation and emission slit widths of 9 nm and 20 nm. Four circular areas were measured per well. Background fluorescence was determined from wells without cells and subtracted from all measurements. Mean ratios were obtained from triplicate wells for calibration experiments and from duplicate wells for H₂O₂ stimulation experiments, based on the average ratios across the four areas.

For calibration experiments, a mixture of saponin (final concentration 40 µg/mL) and ADPR (final concentration 10 nM to 1 mM) in Ca²⁺ measurement buffer was added to the wells. Fluorescence ratios of the ADPR biosensor and the mutant control were determined after 3 min of measurement. Calibration data were plotted against ADPR concentration and fitted to a four-parameter sigmoidal model using GraphPad Prism (v10.5.0).

For experiments in intact cells, H₂O₂ (Sigma-Aldrich, Germany) was added at final concentrations of 10, 50, 100, 500, or 1000 µM after 165 s of baseline measurement; Ca²⁺ measurement buffer served as a control. For inhibition of ADPR production, cells were pre-incubated with the PARP-1 inhibitor PJ34 (10 µM; EMD Millipore, Merck, Darmstadt, Germany) for 20 min at 37 °C prior to fluorescence recording.

### Determination of intracellular ADPR

HEK293T cells were seeded in 6-well tissue culture test plates (TRP) and maintained in Dulbecco’s Modified Eagle’s Medium (DMEM, Sigma-Aldrich) supplemented with 10% fetal bovine serum (FBS, Biowest) at 37 °C in a 5% CO₂ atmosphere. When cultures reached WT 6.2 × 10⁶ cells/well; H₂O₂ 6.4 × 10⁶ cells/well, H₂O₂ was added to test plates to a final concentration of 100 µM and incubated for 3 min. The medium was then aspirated, and cells were rapidly washed with 10 mM ammonium acetate (4°C), placed on dry ice, and quenched with 500 µL methanol (80%) chilled at -80°C containing 1 µM 2′/3′-O-acetyl-ADPR-d₃ internal standard. After 20 min at -80°C, cells were scraped on dry ice and extracted with an additional 500 µL methanol (80%). Lysates were vortexed (5 min, 4°C) and centrifuged (14,000 × g, 10 min, 4°C), and supernatants were analysed by LC-MS/MS. A medium-only blank (250 µL growth medium) was processed in parallel. ADP-ribose was quantified by LC-MS/MS using an Agilent Infinity II 1290 HPLC (G7116B, bio-inert) coupled to a SCIEX QTRAP 6500+ triple quadrupole mass spectrometer in negative electrospray ionization mode. Separation was performed on an Atlantis Premier BEH Z-HILIC column (2.1 × 100 mm, 1.7 µm; Waters) with mobile phase A (H₂O/ACN 90:10, 10 mM ammonium acetate, pH 9) and B (ACN/H₂O 90:10, 10 mM ammonium acetate, pH 9) at 0.25 mL/min using the following gradient: 5% A (0-2 min), 40% A (14.5-16 min), and re-equilibration to 5% A (16.5-20 min). ADP-ribose was quantified by MRM using optimized transitions (m/z 558.050 to 346.118 (quantifier), 78.930) with 2′/3′-O-acetyl-ADPR-d₃ as a structural-analogue internal standard (m/z 603.104 to 346.097 (quantifier), 78.919). Absolute quantification was performed using a 7-point calibration curve (0.015-100 µM, R² = 0.986) based on analyte-to-internal-standard peak area ratios. Intracellular concentrations were calculated assuming a cell volume of 1.89 pL per adherent HEK293 cell (Jang et al., 2022). Medium-only concentrations were calculated assuming total extraction volume.

### HPLC analysis of nucleotides

Stability of 8-Br-cADPR under nDSF conditions was assessed by reversed phase (RP)-HPLC. Reaction products were analysed using a 250 mm x 4.6 mm C18 Multohyp BDS column (5 µm particle size, CS-Chromatographie Service GmbH) in conjunction with a C18 Multohyp BDS guard column 17 mm x 4.6 mm (5µm particle size, CS-Chromatographie Service GmbH). The mobile phase consisted of HPLC buffer A (20 mM KH_2_PO_4_, 5 mM tetrabutylammonium phosphate monobasic solution, pH 6.0) and buffer B (50% buffer A, 50% methanol). To elute the nucleotides from the column the methanol content in the mobile phase (flow rate of 0.8 ml/min) was increased over time (0.0 min [30% buffer B], 3.0 min [30% buffer B], 11.0 min [62.5% buffer B], 25.0 min [100% buffer B], 27.0 min [100% buffer B], 38.0 min [30% buffer A], 43.0 min [30% buffer A]). The adenine nucleotides were detected by their light absorption at 260 nm using a diode-array detector (DAD, Agilent Technologies). For quantification standards with different concentrations of commercial 8-Br-ADPR (#B 051, Biolog) and 8-Br-cADPR (#B 065, Biolog) were used.

### Statistical analysis

Data sets were assembled in Excel (Microsoft Office Standard 2016), whereas statistical analysis and visualization were performed in GraphPad Prism (v10.5.0). Log-transformed current amplitudes from patch-clamp experiments were analyzed using a two-way ANOVA (genotype vs. nucleotide). A significance level (α) of 0.05 was used for all statistical tests. Data analysis and visualization of the measurements obtained with the ADPR biosensor were performed using Python (v3.14), with Matplotlib, NumPy, and SciPy in Jupyter notebooks.

## Author contributions

T.K. and T.T. expressed and purified recombinant proteins. T.K., T.T., and L.D. performed nDSF and ITC experiments. T.S, S.E. and R.F. performed the electrophysiology experiments. F.K. performed the Quick Change mutagenesis to generate the N1487A construct used for the electrophysiology experiments. V.N., S.G. and X.A.C. provided biosensor measurement expertise and the ADPR biosensor cell line, T.S. and R.F. performed the in situ calibration and the stimulation experiments with the biosensor cell lines. V.P. and M.Z. performed the metabolomics experiments. A.H.G. contributed to the study design by proposing the implementation of the biosensor approach. M.G.A. conceived and supervised the project together with H.T. and R.F. .T.K., M.G.A. and R.F. wrote the manuscript with input from all authors.

## Acknowledgements

We acknowledge access to the Sample Preparation and Characterization (SPC) Facility of EMBL, Hamburg. This work was supported by the Deutsche Forschungsgemeinschaft (DFG) (SFB1328, project A05 to RF and MGA and project A01 to AHG), as well as the NIH (R01CA272490, R35GM152218 to XAC).

## Notes

The authors declare no competing financial interest. S.G and X.A.C disclose that they are inventors on a US patent application filed by The University of Texas at Austin covering the ADPR biosensor technology.

## Footnotes

The abbreviations used are:

dADPR: 2’-deoxy-adenosine diphosphate ribose
ADPR: adenosine diphosphate ribose
CI95: asymmetric 95% confidence intervals
dr: *Danio rerio* (zebrafish) DMEM Dulbecco’s Modified Eagle Medium
EDTA: ethylenediaminetetraacetic acid
EM: electron microscopy
FBS: fetal bovine serum
hs: *Homo sapiens* (human)
IMAC: immobilized metal affinity chromatography
IPTG: isopropyl β-D-1-thiogalactopyranoside
ITC: isothermal titration calorimetry
MHR: TRPM homology region
MOI: multiplicity of infection
NAD: nicotinamide adenine dinucleotide
Ni-NTA: Ni-nitrilotriacetic acid
NMDG: *N*-methyl-D-glucamine
PARG: poly(ADP-ribose) glycohydrolase
PARP: poly(ADP-ribose) polymerase
PEI: polyethylenimine
ROS: reactive oxygen species
SDS-PAGE: sodium dodecyl sulphate-polyacrylamide gel electrophoresis
SD: standard deviation
SEM: standard error of the mean
TCEP: tris(2-carboxyethyl)phosphine
TEV: tobacco etch virus
TRPM: transient receptor potential melastatin
WT: wild type.

