## Supplementary Information for "Domain-Specific Agonist Binding Affinities Explain Structural and Functional Regulation of TRPM2"

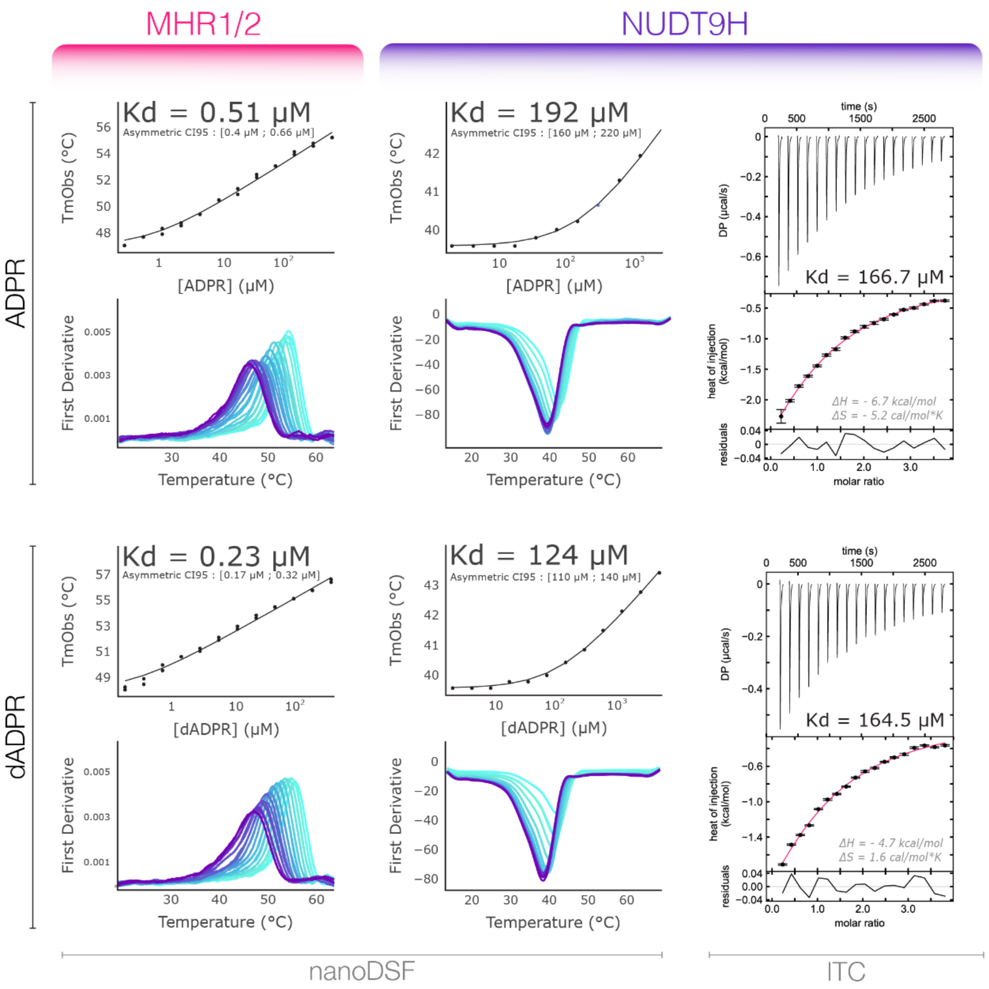


**Supplementary Figure 1. Binding affinities of ADPR and dADPR to TRPM2 domains measured by nDSF and ITC.** Dissociation constants ($K_d$s) were determined for the MHR1/2 and NUDT9H domains separately. nDSF measurements revealed that the $K_d$ for MHR1/2 (ADPR = 0.51 μM CI95 [0.44 μM; 0.66 μM] and dADPR = 0.23 µM) CI95 [0.17 µM; 0.32 µM] is nearly three orders of magnitude lower than that for NUDT9H (ADPR = 192 µM CI95 [160 µM; 220 µM], dADPR = 124 µM CI95 [110 µM; 140 µM]), indicating much stronger ligand binding. For the NUDT9H domain, $K_d$ values were additionally confirmed by isothermal titration calorimetry (ITC) (ADPR = 166.7 µM, dADPR = 164.5 µM). In nDSF experiments, where the peak corresponds to the observed melting temperature ($T_{mObs}$). $T_{mObs}$ against ligand concentration depicts the ligand-induced stabilization of the protein; a concentration-dependent shift in $T_{mObs}$ reflects ligand binding and is used to calculate the dissociation constant.

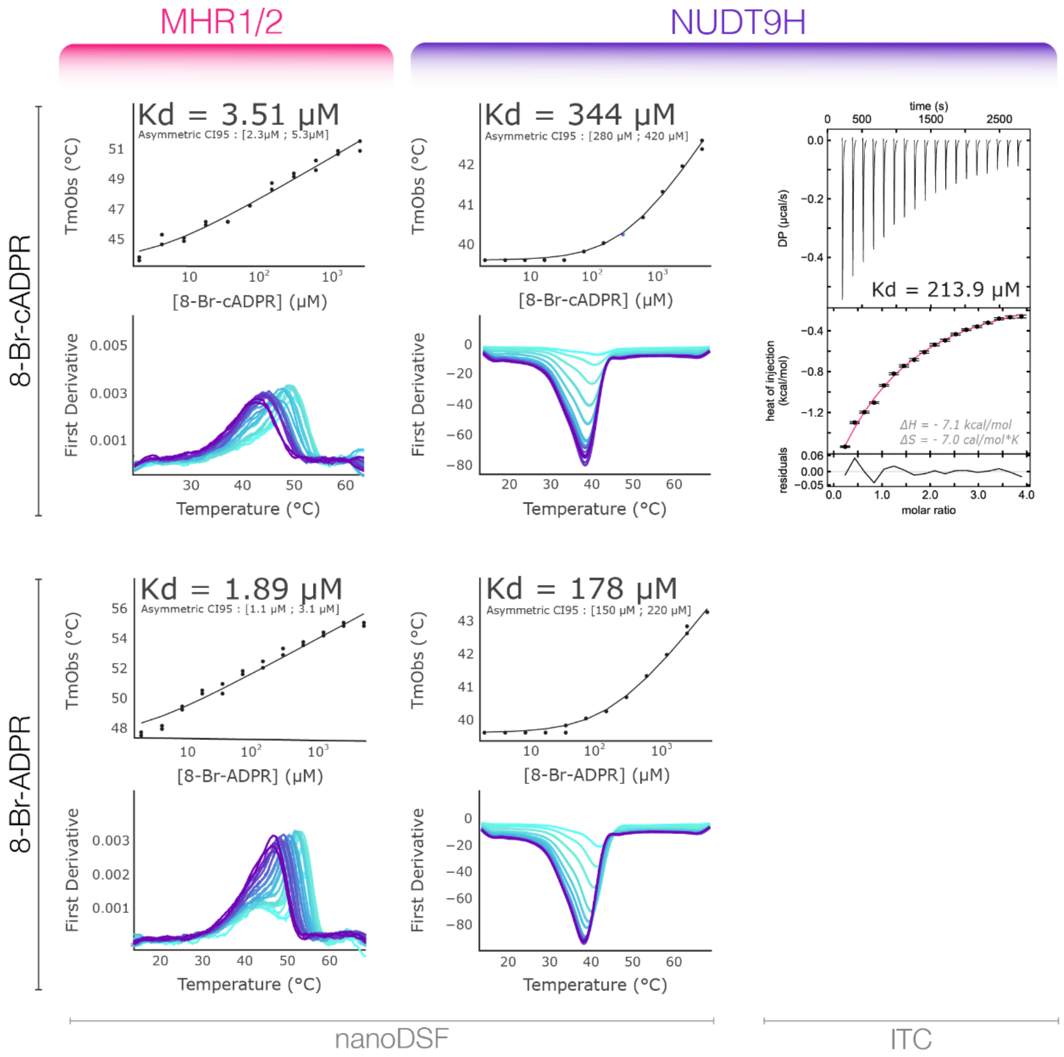


**Supplementary Figure 2. Binding affinities of 8-Br-cADPR and 8-Br-ADPR to TRPM2 domains measured by nDSF and ITC.** Dissociation constants ($K_d$s) were determined for the MHR1/2 and NUDT9H domains separately. For the NUDT9H domain, $K_d$ values were additionally confirmed by isothermal titration calorimetry (ITC).

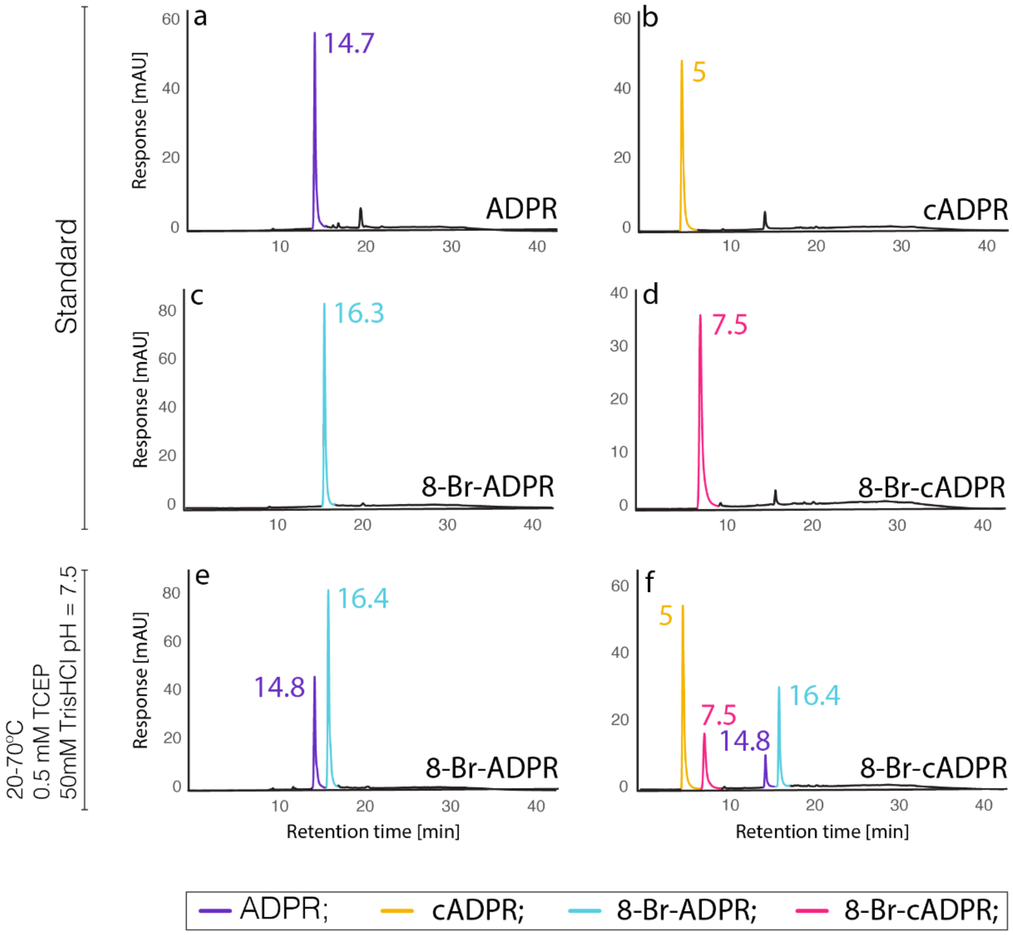


**Supplementary Figure 3. HPLC analysis of degradation of 8-brominated nucleotides. a-d -** Standards: a - ADPR, b - cADPR, c - 8-Br-ADPR, d - 8-Br-cADPR; e–f - Chromatograms of 8-Br-ADPR and 8-Br-cADPR after treatment under nDSF conditions (gradual heating from 20 °C to 70 °C in 50 mM Tris-HCl, pH 7.5, with 0.5 mM TCEP). Under nDSF conditions 8-Br-ADPR degrades to ADPR. 8-Br-cADPR degrades not only into 8-Br-ADPR but also into cADPR and ADPR. Peaks are color-coded within a single chromatogram (chromatograms are representative of three independent experiments).

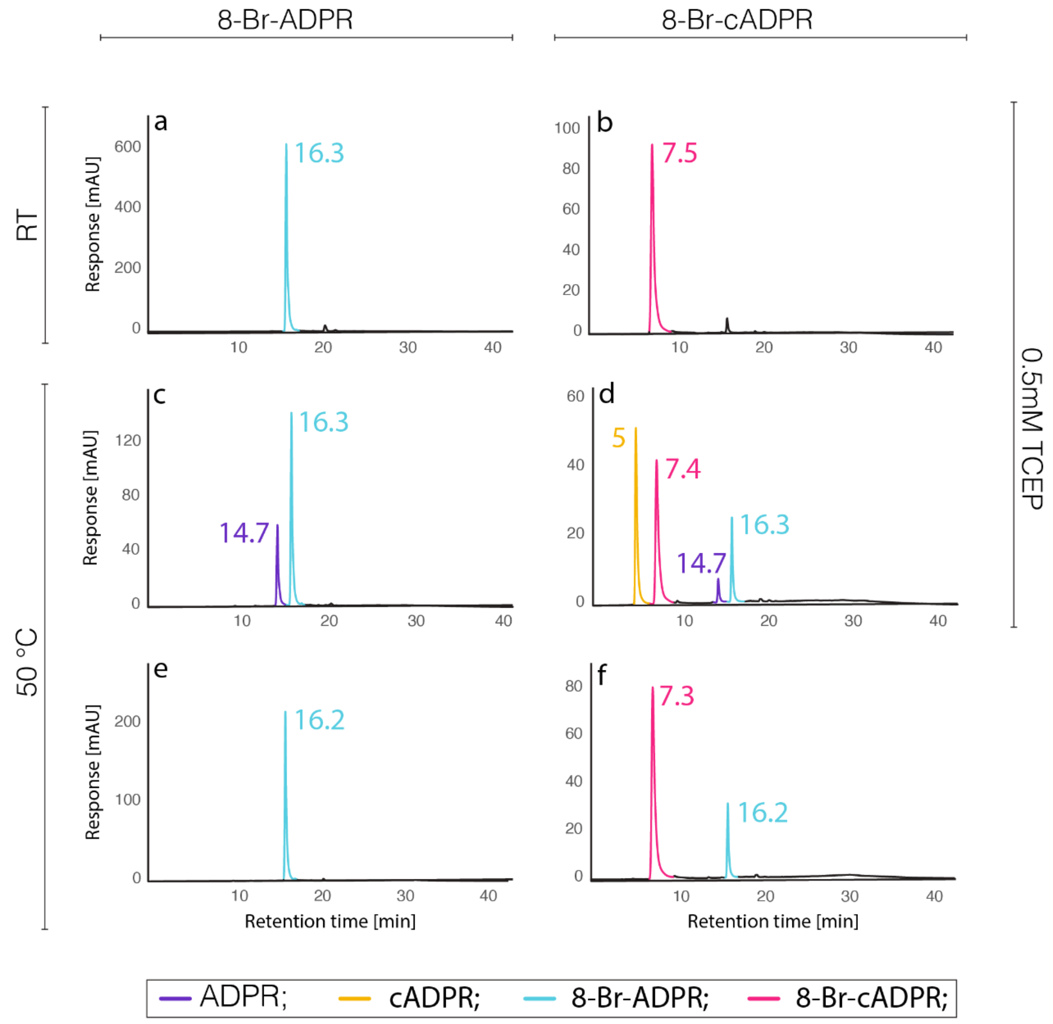


**Supplementary Figure 4. HPLC analysis of the impact of temperature and TCEP and temperature on 8-brominated-nucleotides.** Comparison of the stability of 8-Br-cADPR and 8-Br-ADPR at room temperature (a, b) and 50 °C, in the presence (c, d) or absence of TCEP (e, f). TCEP or heat alone had only minor effects, whereas their combination strongly promoted nucleotide breakdown. Peaks are color-coded within a single chromatogram. (Chromatograms are representative of three independent experiments)

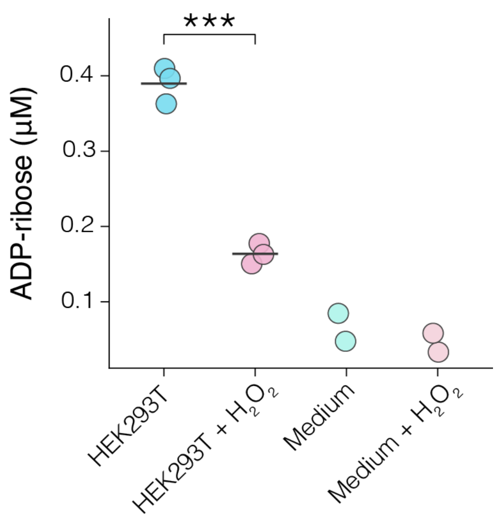


**Supplementary Figure 5. Intracellular ADPR concentration.** Intracellular ADP-ribose concentrations were determined by LC-MS/MS in untreated wild-type (HEK293T) cells and cells exposed to $H_2O_2$ (HEK293T + $H_2O_2$). Each dot represents an independent biological replicate and horizontal bars indicate the mean (n = 3). $H_2O_2$ treatment significantly reduced cellular ADP-ribose levels compared to WT (two-tailed unpaired Welch's t-test, $p$ = 6.03 × $10^{-4}$). Medium and Medium + $H_2O_2$ represent medium-only blanks processed in parallel and calculated assuming total extraction volume.

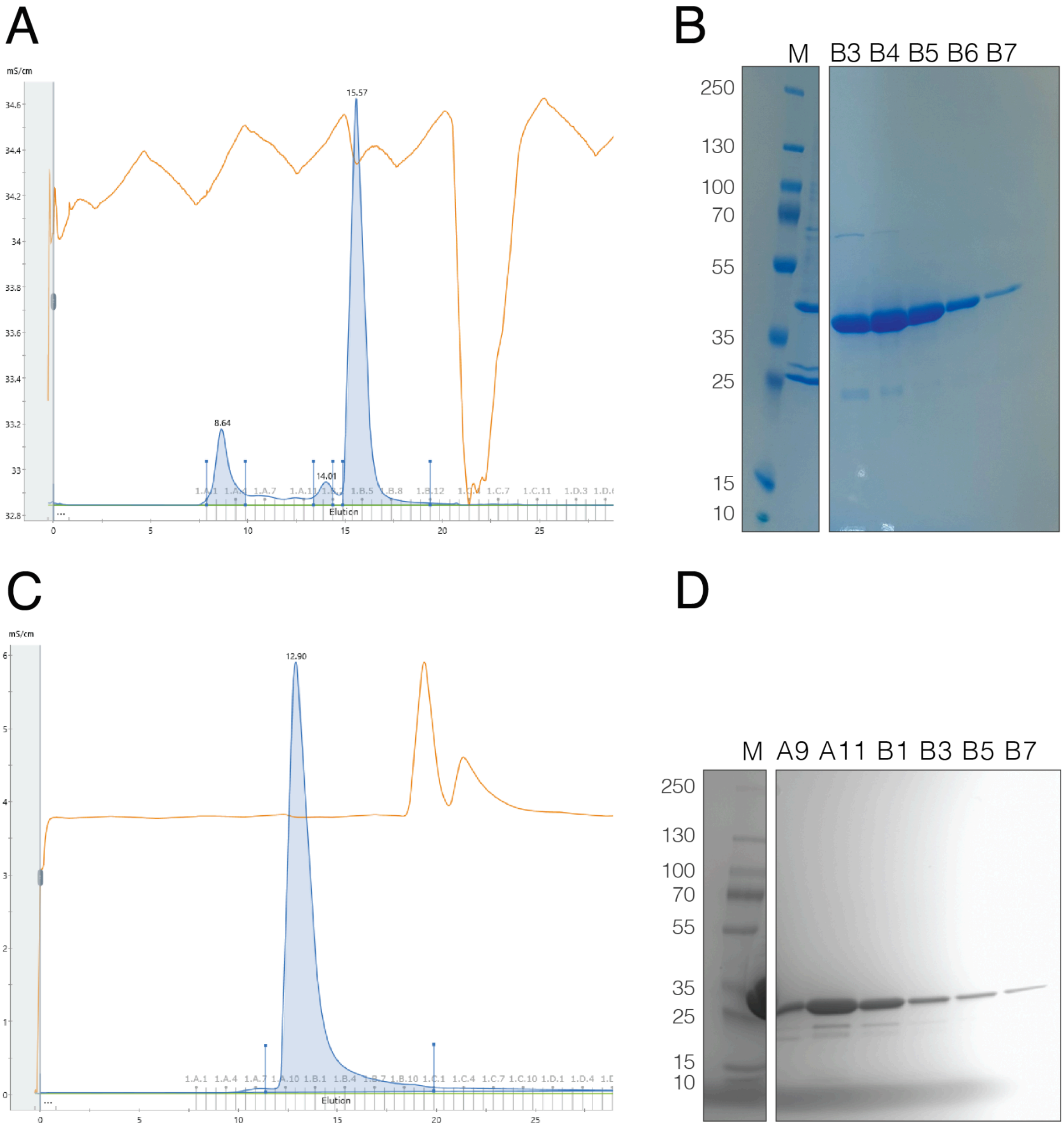


**Supplementary Figure 6. Purification of MHR1/2 and NUDT9H.** SEC and SDS page of MHR1/2 purification (A,B) and NUDT9H (C,D).
